# Multimodal protein representations across human brain cell types

**DOI:** 10.64898/2026.11.10.765762

**Authors:** Amy Dong, Michelle M. Li, Kerr Ding, Sumaiya Nazeen, Ayush Noori, Yepeng Huang, Shanghua Gao, Dylan Ritter, Jalwa Afroz, Alexander B. H. Henderson, Lorenz Studer, Yunan Luo, Vikram Khurana, Giovanni Parmigiani, Marinka Zitnik

**Affiliations:** Department of Biomedical Informatics, Harvard Medical School, Boston, MA, USA; Department of Biomedical Engineering, College of Engineering, Carnegie Mellon University, Pittsburgh, PA, USA; School of Computational Science and Engineering, Georgia Institute of Technology, Atlanta, GA, USA; Aligning Science Across Parkinson’s (ASAP) Collaborative Research Network, Chevy Chase, MD, USA; Department of Neurology, Brigham and Women’s Hospital, Boston, MA, USA; Department of Engineering Science, University of Oxford, Oxford, UK; The Center for Stem Cell Biology, Memorial Sloan Kettering Cancer Center, New York, NY, USA; Broad Institute of MIT and Harvard, Cambridge, MA, USA; Harvard Stem Cell Institute, Cambridge, MA, USA; Department of Data Science, Dana-Farber Cancer Institute, Boston, MA, USA; Department of Biostatistics, Harvard T.H. Chan School of Public Health, Boston, MA, USA; Kempner Institute for the Study of Natural and Artificial Intelligence, Harvard University, MA, USA; Harvard Data Science Initiative, Cambridge, MA, USA

**Author notes:** Equal contribution (Amy Dong, Michelle M. Li). SEMPER website: https://zitniklab.hms.harvard.edu/SEMPER. SEMPER code: https://github.com/mims-harvard/SEMPER.

## Abstract

The central nervous system (CNS) comprises diverse cell types whose molecular context shapes the roles of the proteins they express. Computational models offer a scalable way to study the roles of proteins across human brain cell types, with protein language models and interactome- or single-cell-based models capturing complementary information encoded in protein sequences and cellular network contexts. Human brain tissue is hard to access, so brain cell types make up a small fraction of the atlases single-cell models train on and of the benchmarks used for their evaluation. To capture these complementary sources of information in the CNS, we introduce SEMPER, a multimodal representation learning model that aligns protein language model representations with protein-protein interaction networks from 442 cell types of the adult human brain, giving each protein a sequence-informed representation in every cell type in which its gene is expressed. We show that SEMPER representations predict cell type-specific protein complexes and predict genes essential to the survival of dopaminergic neurons, measured in a genome-wide CRISPR screen. In zero-shot retrieval, SEMPER recovers clinically pursued drug targets for Parkinson’s disease, Alzheimer’s disease, multiple sclerosis, Huntington’s disease and amyotrophic lateral sclerosis at higher rates than cell type-specific models. To further improve target recovery, we combine SEMPER with a large language model that reasons over protein functional annotations. The combination recovers more targets, with SEMPER providing experimentally derived cellular context that complements literature-based reasoning of the large language model.

## Main

The central nervous system (CNS) comprises diverse cell types in which the same protein can have distinct biological roles. A protein’s sequence encodes intrinsic properties that shape these roles, and the cell type contributes its expression, its interaction partners, and the molecular processes active in that context [1–5]. Parkinson’s disease illustrates how these complementary sources of information act together. *α*-Synuclein, encoded by *SNCA*, is expressed across many CNS cell types, yet the disease is characterized by the preferential loss of dopaminergic neurons in the substantia nigra [6, 7]. Pathogenic *SNCA* variants alter the structure and aggregation of *α*-synuclein [8], while the cell types that express the protein differ in expression, interaction partners, and active pathways. Huntington’s disease follows the same pattern, where an expanded CAG repeat in *HTT* lengthens the polyglutamine tract of huntingtin [9] and the effects of the mutant protein depend on its interaction partners [10, 11]. Characterizing the roles of proteins in CNS biology and disease therefore depends on their sequence-encoded properties and on their cellular context.

Protein language models learn biochemical and evolutionary information from protein sequences [12–19]. Trained on hundreds of millions of sequences across the tree of life, these models represent proteins by the constraints that evolution has placed on their residues, which are informative of their structure, their biochemical properties, and the effects of variants in their sequences. Network-based models instead learn protein representations from molecular interactions, restricted to proteins whose genes are expressed in a given cell type [20–24]. Protein representations then depend on the interaction partners present in that cell type, so the same protein has a different representation in every cell type with a different interaction neighborhood. Single-cell foundation models use atlas-scale transcriptomic datasets to learn representations across cell type contexts [25–33]. These datasets draw on tissues from across the body, and human brain tissue is harder to obtain than organs accessible through blood and biopsy, which are sampled from living donors as a matter of routine. Brain cell types therefore account for a small share of the data these models train on and of the datasets used for their evaluation, and models of single-cell data generalize poorly to cell types with sparse coverage in their training data [34, 35]. In the CNS, single-cell and network-based approaches have prioritized cell type-specific genes and therapeutic targets in neurodegenerative disease and have reconstructed disease-relevant molecular networks [36, 37]. Each of these approaches learns from one source of information, so proteins are represented either by their sequences or by interactions within their cell type of expression. A model that aligns the two sources, and that learns from CNS cell types alone, would represent proteins within each cell type while retaining the properties encoded in their sequences, and would support more accurate prediction of cell type-specific protein function and therapeutic relevance.

Here, we introduce SEMPER, a multimodal representation learning model that aligns a sequence-based protein language model with a graph transformer of protein interactions across 442 cell type-specific protein-protein interaction networks of the adult human brain. SEMPER projects the outputs of the two encoders into a shared latent space, where the sequence information of a protein is paired with its interaction environment in each cell type, and combines them into a cell type-resolved representation for every protein in each cell type of expression. Proteins have different representations across cell types while retaining the information encoded in their sequences, which distinguishes SEMPER from contextual models trained on interaction networks alone.

Within SEMPER’s latent space, proteins are organized by cell type and biological function, and their representations predict cell type-specific CNS protein complexes more accurately than existing models under four complementary complex definitions. In zero-shot retrieval, without task-specific training, SEMPER recovers clinically pursued therapeutic targets for Parkinson’s disease, Alzheimer’s disease, multiple sclerosis, Huntington’s disease, and amyotrophic lateral sclerosis at higher rates than existing cell type-specific models. Recovery varies across cell types within each disease, so the representations indicate the cellular contexts in which the targets of a disease are most readily recovered. SEMPER’s representations also provide experimentally grounded information that complements the literature-based knowledge of large language models. SEMPER predicts experimentally identified essential genes in dopaminergic neurons more accurately than a large language model alone, and a large language model that compares SEMPER’s candidate genes using retrieved Gene Ontology annotations further improves recovery of clinically pursued targets. Together, these results establish cell type-resolved protein representations as an approach to study protein function across CNS cell types and to ground language model reasoning in experimental measurements.

## Results

### Cell type-resolved multimodal protein representations

SEMPER learns protein representations by combining protein sequence, cell type-specific protein-protein interactions, and the hierarchical organization of CNS cell types (Figure 1a; Methods 2). We construct cell type-specific protein–protein interaction (PPI) networks for 442 CNS cell types by intersecting a single-nucleus RNA sequencing atlas of the adult human brain, comprising ∼3.37 million cells from 112 anatomical locations [38], with a cell type-agnostic interactome [39], yielding more than 1.25 million protein nodes across the 442 cell type networks (Methods 1.1). A metagraph with 2,316 edges between 599 cell types and tissues encodes the hierarchical and anatomical relationships among these cell types. Each of the 15,487 unique proteins in these networks has one reference sequence and appears in an average of 81 cell type-specific networks (Methods 1.1). By integrating these networks with protein sequences, SEMPER learns a multimodal embedding space of proteins based on physical interactions, cell type-specific expression, anatomy, and protein sequences.

**Figure 1:**
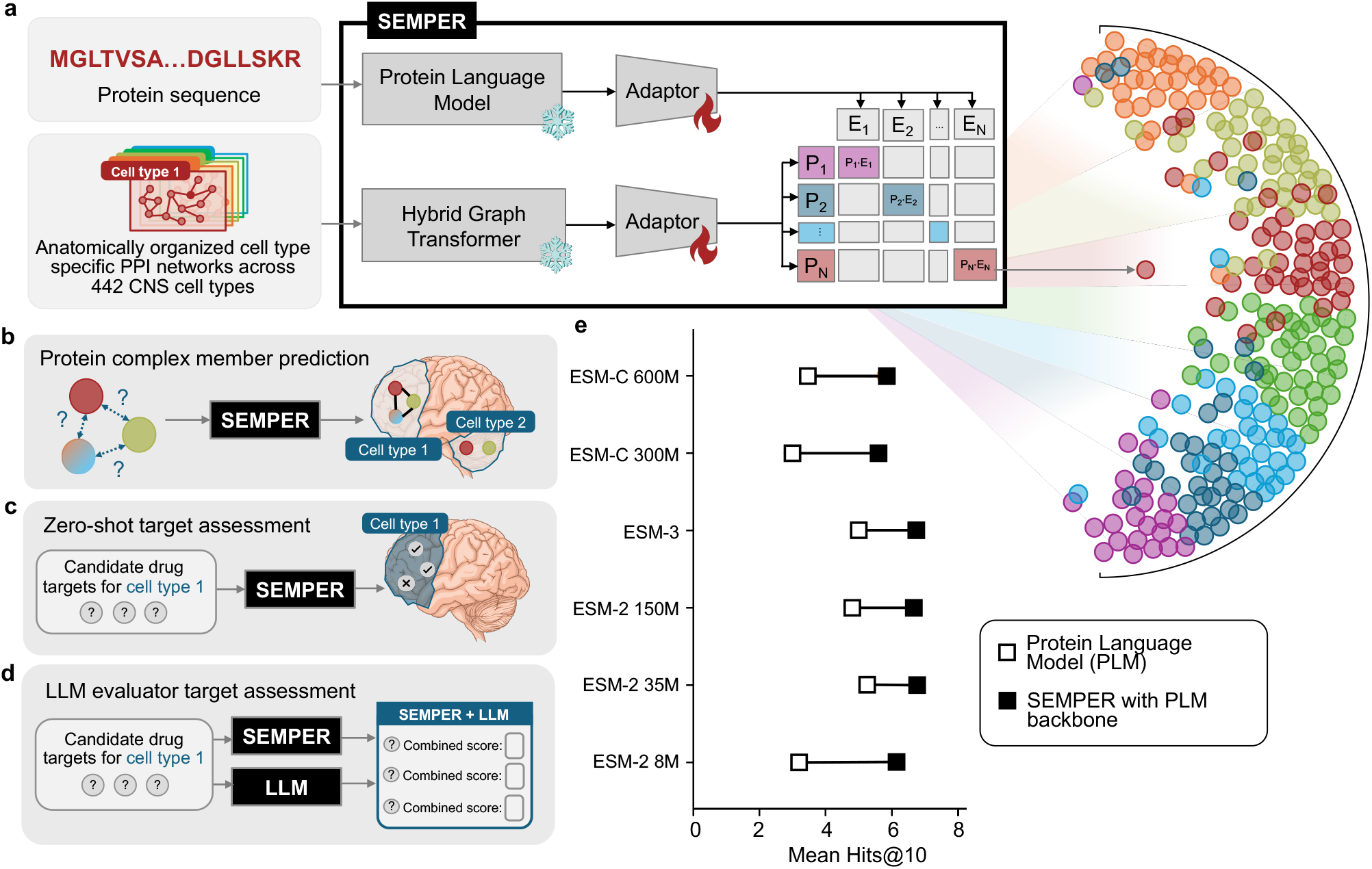
Overview of SEMPER. **(a)** SEMPER is a multimodal model trained on 442 cell type-specific protein-protein interaction (PPI) networks constructed by intersecting a protein interactome with a single-cell transcriptomic atlas of the adult human central nervous system (CNS), protein sequences encoded by a protein language model trained on hundreds of millions of protein sequences, and a hierarchical cell-tissue anatomical metagraph of the CNS. A protein language model encodes protein sequences, and a hybrid graph transformer encodes the context of each protein within the cell type-specific PPI networks and the CNS metagraph. Trainable adaptors project the outputs of the two encoders into a shared latent space, where a symmetric InfoNCE contrastive loss aligns the sequence representation of each protein with its cell type-specific context representations. **(b-d)** Applications of SEMPER. **(b)** SEMPER predicts whether a set of proteins assembles into a complex in a given CNS cell type. **(c)** In zero-shot retrieval, SEMPER retrieves candidate drug targets within a given cell type, ranking proteins by their similarity to a small set of known targets. **(d)** A large language model scores and reorders SEMPER’s candidate drug targets. **(e)** Mean Hits@10 for zero-shot retrieval of clinically pursued targets curated across neurodegenerative diseases, comparing each protein language model alone (Protein Language Model, open squares) with SEMPER built on that protein language model (SEMPER with PLM backbone, filled squares). The protein language models span three families, grouped by family and ordered by parameter count within each family (ESM-2 8M, 35M, 150M; ESM-3 1.4B; ESM-C 300M, 600M).

SEMPER uses separate encoders for protein sequence and network interaction cell type context, followed by trainable adaptors that map both representations into a shared latent space. The graph encoder is a 1.3-billion-parameter hybrid graph transformer [40] trained on the 442 cell type-specific PPI networks and metagraph, producing a unique protein representation for each cell type context (Methods 2.1.1). The sequence encoder is a protein language model (PLM) that outputs a cell type-agnostic protein embedding based on its reference protein sequence (Methods 2.1.3). Here, we leverage ESM models [12, 13] as the sequence encoder, although SEMPER can use any PLM. SEMPER projects the output of each frozen encoder through a trainable adaptor and aligns the projections in a shared latent space with a symmetric InfoNCE objective [41, 42] to produce multimodal protein representations (Methods 2.2). For a protein expressed in multiple cell type contexts, SEMPER learns an embedding that is anchored to its protein sequence and varies with its cell type-specific interactions.

We use SEMPER to interrogate cell type-specific protein behavior across several applications (Figure 1b-d). Given a set of proteins, SEMPER predicts whether they assemble into a complex in a particular cell type (Figure 1b). For neurodegenerative diseases, SEMPER retrieves candidate therapeutic targets specific to each cell type using a small set of known targets as queries (Figure 1c). Representations carry cell type-specific, experimentally-derived information that can complement the reasoning ability of LLMs (Figure 1d). Across six protein language model backbones spanning three ESM families, SEMPER recovers more clinically pursued targets than the backbone alone (Figure 1e), showing that aligning sequence with cell type-specific interactions improves on the sequence model it builds on.

### SEMPER’s representations reflect cell type and biological function

SEMPER represents each protein by aligning complementary modalities in a shared latent space. Protein sequences contain information about protein function, the cell type-specific protein interaction networks reflect the differences in gene expression and protein interactions across cell types, and the metagraph captures the hierarchical organization of cell types and tissues. We assess whether SEMPER’s multimodal protein representations reflect known biology, particularly by cell type specificity and biological function (Methods 3.1.1).

We hypothesize that SEMPER’s protein representations yield cell type-specific protein representations, such that two proteins within the same cell type share higher similarity than proteins across distinct cell types. Across the 30 CNS cell-type superclusters, we find that intra-cell-type (i.e., proteins from the same cell type) similarity is higher than inter-cell-type (i.e., proteins from different cell types) similarity in every supercluster (Figure 2a; Mann-Whitney U test [43] with Benjamini-Hochberg false-discovery-rate correction [44], adjusted *P* < 0.001 for all comparisons; Supplementary Table 1). This shift holds across neuronal, glial, and non-neuronal lineages, showing that SEMPER retains cell type identity.

**Figure 2:**
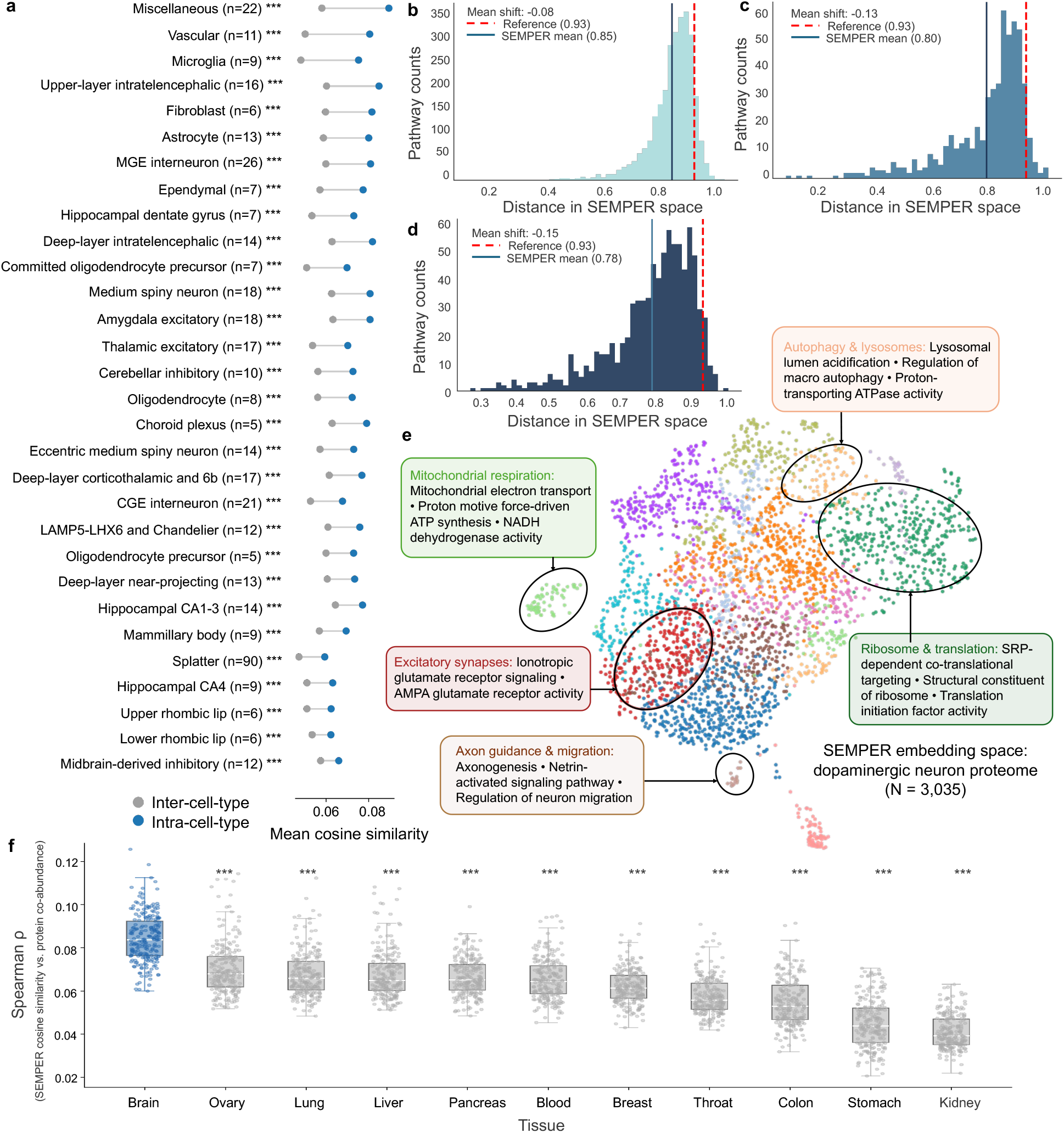
SEMPER organizes proteins by cell type and biological function. **(a)** Mean pairwise cosine similarity of protein pairs sampled within the same cell type (intra-cell-type, blue) compared with protein pairs sampled between that cell type and a different cell type (inter-cell-type, grey), grouped across 30 CNS superclusters. **(b-d)** Distribution of the mean pairwise cosine distance between the SEMPER representations of proteins sharing a Gene Ontology (GO) term, for **(b)** Biological Process, **(c)** Cellular Component, and **(d)** Molecular Function. Solid blue lines mark the mean distance across GO terms in each domain, and dashed red lines mark the mean pairwise distance across all proteins, which serves as the reference distance. **(e)** UMAP projection of the SEMPER representations of the 3,035 proteins in the dopaminergic neuron network. Colors indicate the 15 modules identified by the Leiden algorithm. Selected modules are annotated with their three most enriched GO terms. **(f)** Spearman correlation between the pairwise cosine similarities of SEMPER representations and protein co-abundance scores from the tissue proteomics atlas of Laman Trip et al. [52], computed per cell type across 11 human tissues. We restrict the analysis to cell types in which the fraction of brain-specific protein pairs, defined as pairs with a brain co-abundance score > 0.68 (95th percentile) and a maximum score across all other tissues < 0.5, exceeds the median across all 442 cell types (*n* = 221). Each point represents one cell type.

Across biological process (BP), molecular function (MF), and cellular component (CC) annotations in Gene Ontology [45, 46], proteins sharing a Gene Ontology term lie closer together in the embedding space than randomly sampled proteins, with mean embedding distances lower by Δ_*BP*_ = 0.08 (Fig. 2b), Δ_*CC*_ = 0.13 (Fig. 2c), and Δ_*MF*_ = 0.15 (Fig. 2d). Controlling for the number of proteins per Gene Ontology term and their overlap between terms, we find that same-term proteins remain more compact than average in a permutation test that reassigns embeddings across proteins (≥ 49 SD above the permutation mean; 2,000 permutations; *p*-value < 5 × 10^−4^ for all three domains; Methods 3.1.2), suggesting that SEMPER’s protein representations capture biological function.

We examine dopaminergic neurons, which support voluntary movement and cognition and whose loss underlies Parkinson’s disease [47–49]. Representations of the proteins expressed in dopaminergic neurons cluster into 15 functional modules enriched for pathways central to dopaminergic vulnerability in Parkinson’s disease (Figure 2e; Methods 3.1.3; Supplementary Table 2). The annotated modules span distinct cellular machineries, including mitochondrial respiration and autophagy-lysosomal regulation which are two pathways closely tied to dopaminergic vulnerability [50, 51]. Other highlighted modules include ribosomal translation, excitatory glutamatergic synapses, and axon guidance and neuronal migration (Figure 2e). These modules show that SEMPER organizes the proteins of a single cell type into coherent functional programs.

Leveraging orthogonal experimental data unseen during training, we assess whether SEMPER’s embeddings correlate with brain-specific protein co-abundance. We compare SEMPER’s cosine similarities between protein pairs with an independent proteomics atlas of co-abundance spanning 11 human tissues [52]. We find that SEMPER’s cosine similarities between brain-enriched protein pairs correlate with protein co-abundance more strongly in the brain than each of the other 10 tissues for both the cell types enriched for brain-specific protein pairs and across all 442 cell types (median Spearman *ρ* ≥ 0.082; Figure 2f; BH-corrected one-sided Mann-Whitney U, *p*-value < 0.001 for all comparisons; Supplementary Figure 1; Supplementary Table 3; Methods 3.1.4). In SEMPER’s underlying cell type-specific PPI networks, the existing protein-protein edges have higher brain co-abundance scores than absent edges (Supplementary Figure 2). Altogether, these results show that SEMPER’s latent space reflects experimentally-derived co-abundance scores from independent studies.

### SEMPER predicts cell type-specific protein complexes

Biological function emerges from coordinated protein interactions within macromolecular complexes [53–55]. We test whether SEMPER assigns higher embedding similarity to proteins within curated complexes than to proteins in size-matched decoy sets. We evaluate 547 literature-curated assemblies under four cell type-specific complex definitions, established with transcriptional co-expression, structural PPI connectivity, tissue protein co-abundance [52], and subcellular spatial colocalization [56] (Figure 3a; Methods 1.4; Supplementary Note 1; Supplementary Table 4). We score each candidate complex by the mean pairwise similarity of its members’ embeddings and use the area under the precision-recall curve to measure how well each model separates true complexes from decoys (Methods 3.2). For each definition, we evaluate all models on an identical set of complexes and cell types.

**Figure 3:**
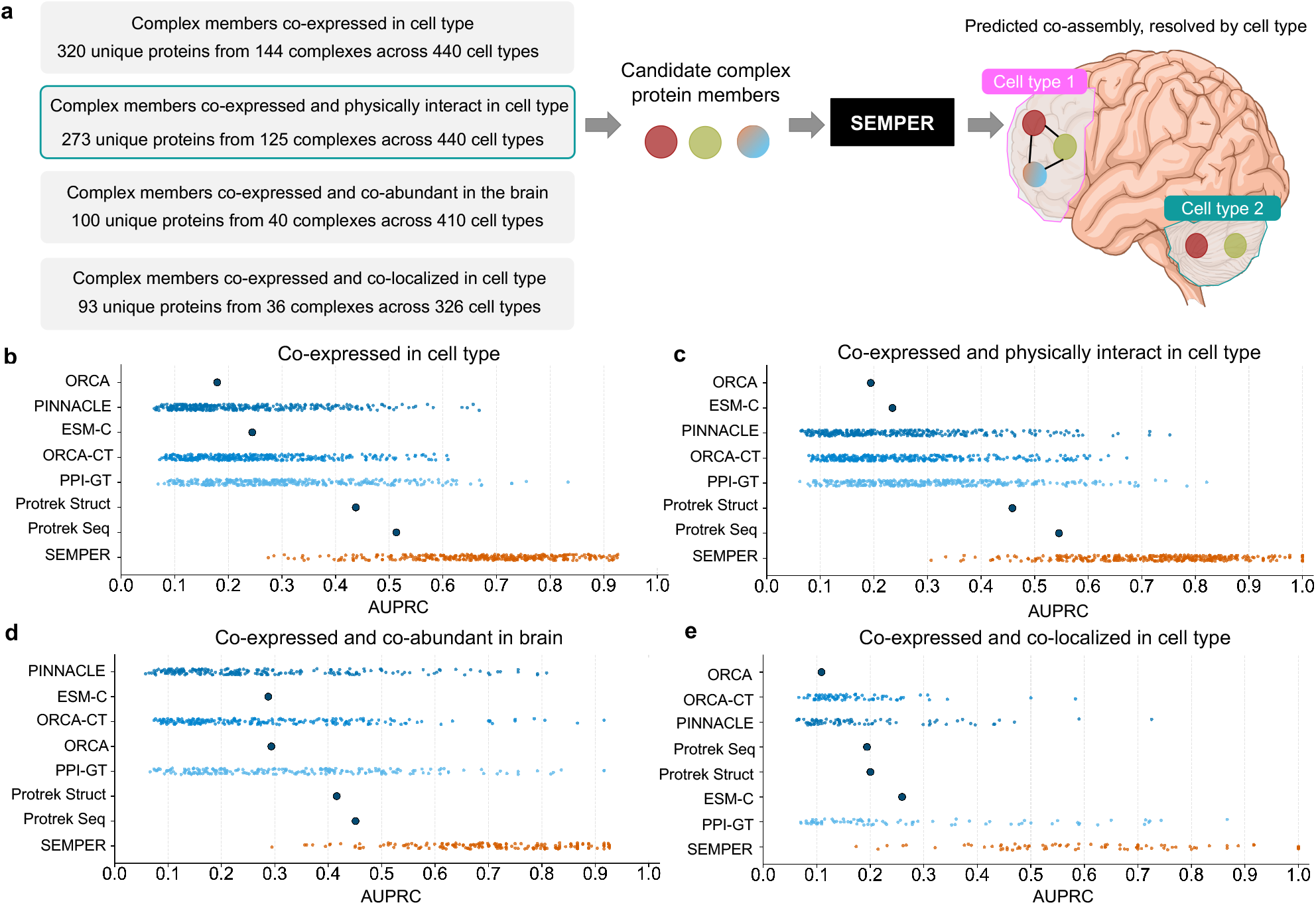
Zero-shot prediction of protein complexes across central nervous system cell types. **(a)** Evaluation framework. Four operational definitions of cell type-specific protein complexes are ordered by strictness, namely co-expression in the cell type, co-expression with physical interaction in the cell type-specific protein–protein interaction (PPI) network, co-expression with co-abundance in brain tissue (co-abundance association score ≥ 0.5 [52]), and co-expression with co-localization to the same subcellular compartment (Human Protein Atlas annotations). For a set of candidate proteins in a given cell type, SEMPER scores the proteins by the mean pairwise similarity of their cell type-specific representations and predicts whether they assemble into a complex. **(b-e)** AUPRC of SEMPER (orange) and existing models (blue) for predicting cell type-specific protein complexes. Each point is a single cell type for the cell type-specific models (SEMPER, PINNACLE, ORCA-CT, PPI-GT) and a single value for the cell type-agnostic models (ORCA, ESM-C, ProTrek Seq, ProTrek Struct). Complexes are defined by **(b)** co-expression in the cell type, **(c)** co-expression and physical interaction in the cell type-specific PPI network, **(d)** co-expression and brain tissue protein co-abundance [52], and **(e)** co-expression and subcellular co-localization.

We compare SEMPER against two groups of models. Cell type-agnostic models assign one unimodal (ESM-C [13] on reference protein sequences and ORCA [57, 58] on cell type-agnostic protein-protein interactions) or multimodal (ProTrek [59], in sequence and structure variants) representation per protein regardless of cell type. Cell type-specific models resolve cell type specificity but use only a single modality (PINNACLE [20], a graph neural network that learns protein representations from cell type-specific PPI networks; ORCA-CT [57, 58], which learns protein signatures from the local graphlet topology of each protein’s interaction neighborhood in those same networks; and PPI-GT, SEMPER’s own graph transformer read out before sequence alignment). We hypothesize that SEMPER’s multimodal, cell type-specific representations identify true complex subunits more effectively by capturing both sequence identity and local cellular context.

SEMPER predicts CNS-specific macromolecular complexes more accurately than existing models across all four complex definitions. Under the transcriptional co-expression definition, SEMPER outperforms all cell type-specific baselines across 349 cell types (all *p*-value < 10^−58^; one-sided Wilcoxon signed-rank test; Methods 3.2; Supplementary Table 5) and SEMPER’s median cell type exceeds the cell type-agnostic baselines ESM-C, ProTrek, and ORCA (Figure 3b). Graph-based models’ performance improves under the structural connectivity definition, likely because their representations encode cell type-specific physical protein-protein interactions. Even so, SEMPER outperforms all cell type-specific baselines across 328 cell types (all *p*-value < 10^−54^; Figure 3c). SEMPER’s advantage persists under the two definitions that draw on evidence independent of the cell type PPI networks. Under the protein co-abundance definition, SEMPER outperforms all cell type-specific baselines across 213 cell types (all *p*-value < 10^−35^; Figure 3d). Lastly, SEMPER performs better than all cell type-specific baselines across 79 cell types (all *p*-value < 10^−12^; Figure 3e) under the spatial colocalization definition. Across all four definitions, SEMPER’s median cell type exceeds every cell type-agnostic baseline, including the strongest, ProTrek-Seq (Figure 3b-e). Notably, SEMPER performs strongly under the latter two definitions despite never seeing these experimentally derived measurements during training.

### SEMPER recovers clinically pursued drug targets

Identifying candidate targets in neurodegeneration benefits from resolving protein function in the cell types where targets are expressed and act [52, 60–64]. We reason that an effective cell type-specific model must retrieve candidate targets across cellular contexts, such as in diseases with cell types vulnerable to pathology.

We evaluate SEMPER on five neurodegenerative diseases that represent different cellular and mechanistic bases underlying their pathology. Parkinson’s disease centers on the loss of dopaminergic neurons and on variants encoded in protein sequence [7, 8, 65]. Huntington’s disease is monogenic and region-selective, with striatal vulnerability despite ubiquitous HTT expression [9, 66–68]. Amyotrophic lateral sclerosis is selective for motor neurons and involves proteostasis [69, 70]. Alzheimer’s disease involves microglial and astrocytic disease-associated states that act non-cell-autonomously [71–74]. Multiple sclerosis, whose clinically pursued targets are largely immunomodulatory and symptomatic, is a mechanistically heterogeneous disease, spanning peripheral immunity, CNS inflammation, demyelination, and neurodegeneration [75–78].

To assess whether SEMPER captures cell type-specific target relevance without task-specific training, we evaluate zero-shot target recovery against clinically pursued neurodegenerative drug targets in clinical trials across all five diseases [79–81] (Methods 1.5). Clinically pursued targets are protein-coding genes targeted by drugs that have completed Phase 2 trials or entered Phase 3 or 4 trials for the disease or its ontological descendants (Methods 1.5). This yields 130 targets for Alzheimer’s disease (73, 48, 9 whose highest phase is Phase II, Phase III, and Phase IV, respectively), 166 for multiple sclerosis (Phase II: 47, Phase III: 56, Phase IV: 63), 86 for Huntington’s disease (Phase II: 18, Phase III: 67, Phase IV: 1), 98 for amyotrophic lateral sclerosis (Phase II: 71, Phase III: 17, Phase IV: 10), and 97 targets for Parkinson’s disease (Phase II: 45, Phase III: 18, Phase IV: 34; Figure 4a; Supplementary Table 6). We compare the targets for Parkinson’s against an independent study that manually curates Parkinson’s disease drug trials from 2015 to 2024 [82]. Of the 117 trials in Phase 2-3 that test an agent with a known protein target, 105 of these (90%) test an agent acting on at least one of our targets, supporting the validity of our pipeline for curating clinically pursued targets (Methods 1.5). For each disease and cell type, we provide models with a small set of known clinically pursued targets, rank all remaining expressed proteins by embedding similarity, and measure the recovery of held-out clinical targets in the top ten (mean Hits@10 across 100 bootstrapped sets; Methods 4).

**Figure 4:**
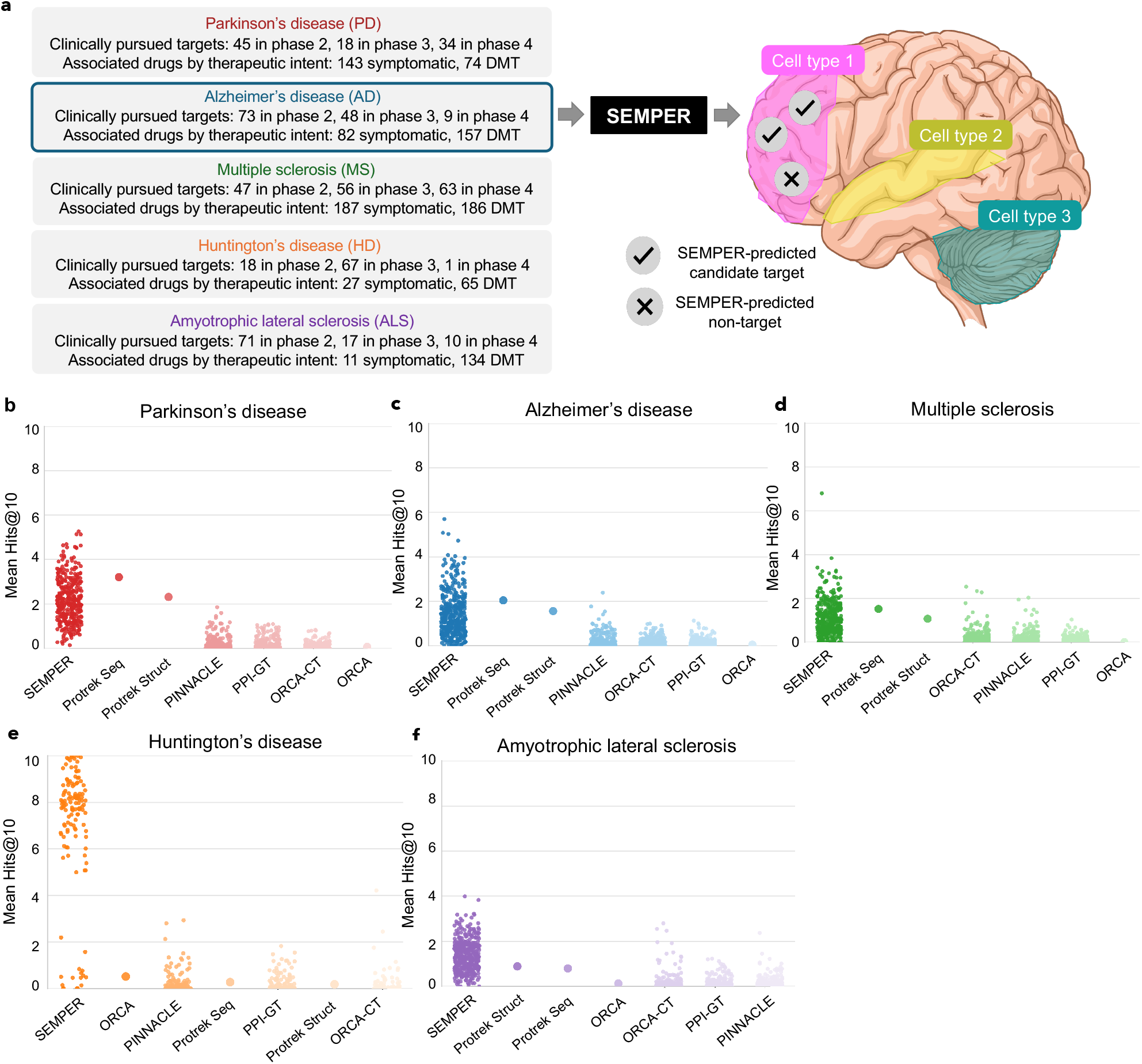
Zero-shot recovery of clinically pursued drug targets for neurodegenerative diseases. **(a)** Evaluation framework for candidate target retrieval. For each of five neurodegenerative diseases (Parkinson’s disease, Alzheimer’s disease, multiple sclerosis, Huntington’s disease, and amyotrophic lateral sclerosis), ground-truth genes are the genes pursued as targets of drug candidates in completed Phase 2 trials or in Phase 3-4 trials of any status. The disease boxes report the number of targets by trial phase and the number of associated drugs classified by therapeutic intent as symptomatic or disease-modifying. For a selected disease, SEMPER receives a randomly sampled subset of these clinically pursued targets as seeds. Within each cell type, SEMPER ranks the remaining proteins in that cell type’s network by their mean similarity to the seed representations in its latent space, and the top-ranked proteins are the SEMPER-predicted cell type-specific candidate targets (checkmarks) and the rest non-targets (crosses). **(b-f)** Number of held-out clinically pursued targets recovered within the top-10 candidates (Hits@10) for **(b)** Parkinson’s disease, **(c)** Alzheimer’s disease, **(d)** multiple sclerosis, **(e)** Huntington’s disease, and **(f)** amyotrophic lateral sclerosis. Each point represents the mean Hits@10 across 100 randomly sampled seed sets within a single cell type for the cell type-specific models (SEMPER, PINNACLE, PPI-GT, ORCA-CT), or a single global mean (solid circles) for the cell type-agnostic models (ORCA, ProTrek Seq, ProTrek Struct).

SEMPER recovers held-out targets at higher rates than all cell type-specific baselines in all five diseases (Figure 4b-f). Against PINNACLE [20], PPI-GT, and ORCA-CT [57, 58], performance gains are significant in every disease (paired one-sided Wilcoxon signed-rank test [83] across cell types, all *p*-value < 10^−21^; Supplementary Table 7). These baselines recover almost no held-out clinically pursued targets (mean Hits@10 ≤ 0.3, s.d. ≤ 0.6, in every disease). Cell type-agnostic models recover targets at rates above zero and are competitive in some diseases. ProTrek-Seq exceeds SEMPER’s mean Hits@10 in Parkinson’s disease (3.2 ± 1.6 versus 2.2 ± 1.0), Alzheimer’s disease (2.0 ± 1.1 versus 1.4 ± 1.0), and multiple sclerosis (1.5 ± 0.9 versus 1.1 ± 0.8), whereas SEMPER is higher in amyotrophic lateral sclerosis (1.4 ± 0.7 versus 0.8 ± 0.5) and Huntington’s disease (7.0 ± 2.8 versus 0.3 ± 0.3). However, cell type-agnostic models represent each protein identically in every cell type, whereas SEMPER represents each candidate within its cellular context, distinguishing the cell types in which targets are most readily recovered and creating a spectrum of cell types with high to low recovery rates. None of the interaction-based models achieves similar performance, including PPI-GT, SEMPER’s own graph transformer before sequence alignment, and SEMPER also exceeds the protein language model it builds on (Fig. 1e), suggesting that the improvement follows from aligning sequence with cell type-specific interaction context.

The range of recovery rates is largest in Huntington’s disease (Figure 4e), where SEMPER reaches a mean Hits@10 of 7.0 of ten across its 129 cell types, and recovers at least six held-out targets in 81.4% of cell types, including 13 cell types exceeding 9.5. In contrast, the strongest baseline, ORCA, reaches a mean of 0.5 and the cell type-specific baselines remain below 0.3. In Parkinson’s disease, SEMPER reaches a mean Hits@10 of 2.2 and recovers three or more targets in 21.4% of cell types, with a maximum of 5.3 (Figure 4b). The Hits@10 are a mean of 1.4 and a maximum of 5.7 in Alzheimer’s disease (Figure 4c), a mean of 1.4 and a maximum of 4.0 in amyotrophic lateral sclerosis (Figure 4f), and a mean of 1.1 and a maximum of 6.8 in multiple sclerosis (Figure 4d). For every disease, SEMPER’s maximum recovery rate across cell types exceeds that of all cell type-specific baselines.

### SEMPER predicts essential genes in dopaminergic neurons

While LLMs capture extensive information regarding protein function, their parametric knowledge covers only evidence described in natural language [84–86]. Because SEMPER learns from experimentally derived data, we evaluate whether SEMPER’s cell type-specific protein representations can complement LLMs in reasoning about CNS [84, 85].

Although CRISPR screens quantify essentiality by targeting gene loci, cellular survival hinges on the functional protein machinery encoded by those genes [87, 88]. We reason that SEMPER’s latent space of proteins can identify whether their corresponding genes are essential to cellular survival by capturing the shared sequence and physical interaction properties of their protein products from cell type-specific networks. We analyze a genome-wide CRISPR screen in dopaminergic neurons of the human midbrain [89] to identify genes required for survival [87] (Methods 5). By using an experimental screen that is not described in natural language, we can assess the contributions of SEMPER and large language models (LLMs) in reasoning about essential genes in dopaminergic neurons. We compare three approaches, in-context prompting of an LLM with a set of example genes [90] (Figure 5a), zero-shot retrieval with SEMPER (Figure 5b), and an LLM that scores and reorders SEMPER’s top 100 candidates (Figure 5c; Methods 6; Supplementary Note 2). The three methods draw on the published literature, the geometry of SEMPER’s representations, and the two in combination, respectively.

**Figure 5:**
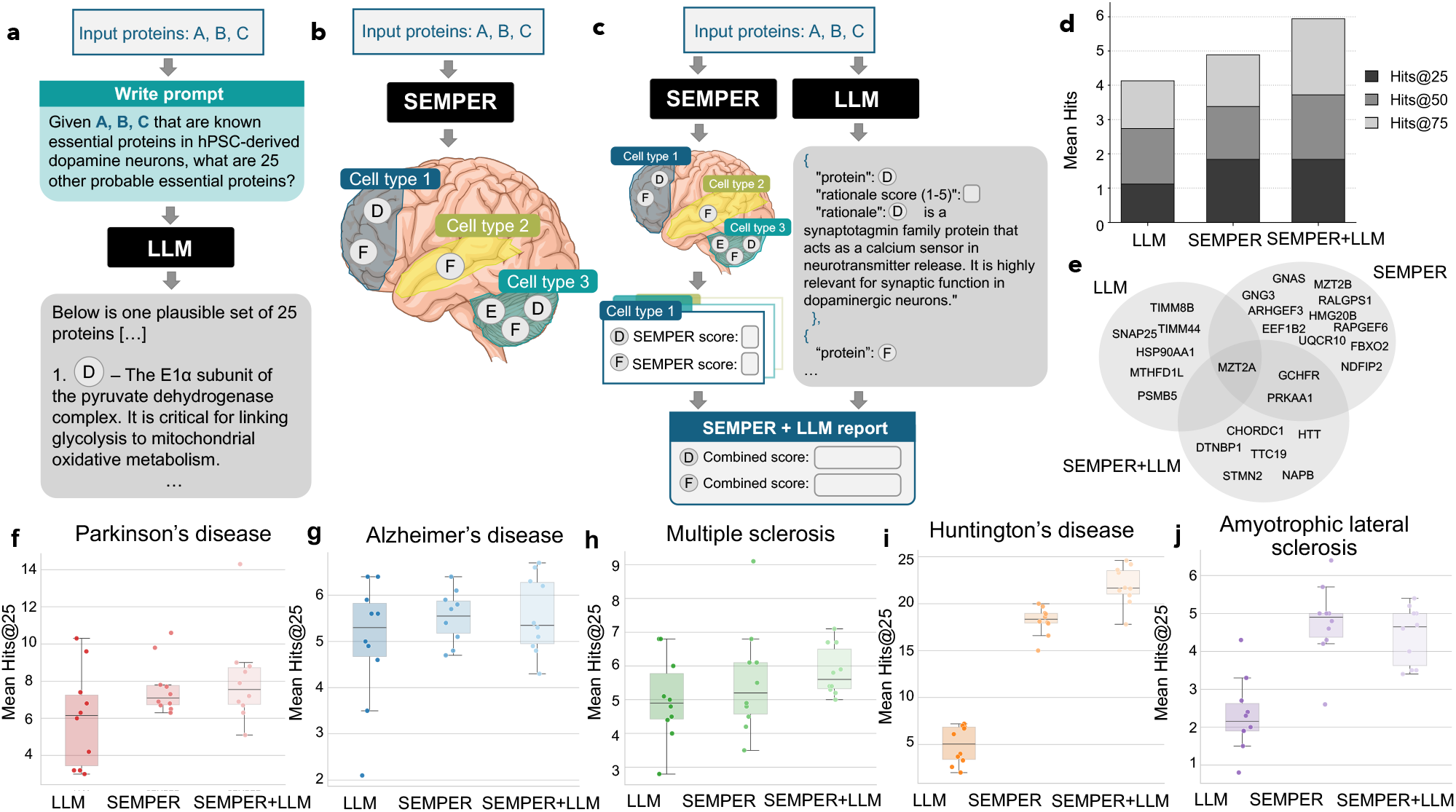
SEMPER and a large language model predict essential genes in dopaminergic neurons and recover clinically pursued drug targets. **(a-c)** Illustration of three zero-shot strategies, labeled LLM-only, SEMPER, and SEMPER+LLM. **(a)** A reasoning large language model (GPT-5.6-sol) receives a set of seed genes and predicts candidate genes from its internal knowledge. **(b)** SEMPER retrieves cell type-specific candidate proteins from the seed proteins. **(c)** A scoring pipeline built on (a) and (b). SEMPER retrieves its top 100 candidate proteins, and a large language model then generates a biological rationale for each protein and scores it on a 1–5 scale. The final ranking averages the large language model scores with SEMPER’s similarity scores. **(d)** Prediction of essential genes in dopaminergic neurons (Hits@25, Hits@50, and Hits@75). **(e)** Venn diagram of the overlap among the three highest-ranked genes in dopaminergic neurons under each strategy. **(f-j)** Recovery of held-out clinically pursued targets within the top 25 candidate genes (Hits@25) for the three strategies (LLM-only, SEMPER, and SEMPER+LLM), evaluated in the ten highest-recovery cell types per disease from Figure 4, for **(f)** Parkinson’s disease, **(g)** Alzheimer’s disease, **(h)** multiple sclerosis, **(i)** Huntington’s disease, and **(j)** amyotrophic lateral sclerosis. Each point is the mean Hits@25 within one cell type. Boxes show the median and interquartile range across the ten cell types, and whiskers extend to the most extreme values within 1.5 times the interquartile range.

On the genome-wide screen, SEMPER predicts essential genes more accurately than the LLM at every threshold (Hits@25 1.8 ± 1.5 versus 1.1 ± 1.2, Hits@50 3.4 ± 1.7 versus 2.7 ± 1.5, Hits@75 4.9 ± 2.2 versus 4.1 ± 1.9; Wilcoxon signed-rank *p*-value ≤ 2.3 × 10^−3^; Figure 5d). This finding is consistent with our hypotheses, as the outputs of the experimental screen are not described in natural language. Reordering SEMPER’s candidates with the LLM yields similar performance on the most stringent thresholds (Hits@25 1.84±1.45 and 1.84±1.15; *p*-value = 0.43) while increasing recovery at broader retrieval depth, raising Hits@50 from 3.38 ± 1.74 to 3.72 ± 1.73 (*p*-value = 0.055) and Hits@75 from 4.89 ± 2.16 to 5.94 ± 1.99 (*p*-value = 7.8 × 10^−5^). The same ordering holds on the 100 genes with the strongest fitness effect (Supplementary Figure 3), with SEMPER again exceeding the LLM at every threshold (Hits@25 0.50 ± 0.56 versus 0.19 ± 0.39, Hits@50 0.93 ± 0.66 versus 0.59 ± 0.60, Hits@75 1.26 ± 0.82 versus 0.85 ± 0.81; *p*-value ≤ 3.3 × 10^−4^), and reordering with the LLM increases recovery further (Hits@25/50/75 of 1.01 ± 0.85/1.67 ± 0.97/2.36 ± 1.09; *p*-value ≤ 8.9 × 10^−7^). Using an LLM to reorder SEMPER candidates allows the LLM to augment SEMPER predictions with complementary literature evidence, improving overall retrieval without injecting genes that are preferentially investigated in the literature or relevant to broad CNS cell types, rather than dopaminergic neurons specifically.

The three approaches rank genes differently (Figure 5e; Supplementary Table 8). SEMPER and the LLM alone produce gene lists with little overlap. Across the top three ranked genes by each method, only *MZT2A*, which encodes a component of the *γ*-tubulin ring complex involved in microtubule nucleation [91], appears among the top three genes of all three approaches. The LLM alone preferentially retrieves well-characterized genes encoding canonical neuronal proteins, including *PSMB5* (a proteasome subunit) [92], *SNAP25* (a presynaptic SNARE protein) [93], and *HSP90AA1* (a molecular chaperone) [94, 95]. SEMPER ranks genes that play a role in mechanistically diverse pathways in dopaminergic neurons, including regulators of heterotrimeric G-protein and small-GTPase signaling (e.g., *GNAS* [96], *GNG3* [97], *ARHGEF3* [98], *RALGPS1* and *RAPGEF6* [99, 100]), mitochondrial respiration (*UQCR10*, encoding a complex III subunit [101, 102]), and neuronal protein trafficking (*FBXO2* [103, 104] and *NDFIP2* [105, 106]). Reordering SEMPER’s candidates with the LLM promotes genes with established neurodegenerative relevance, including *HTT* (whose CAG trinucleotide expansion causes Huntington’s disease) [107, 108], *STMN2* (encoding a neuron-enriched microtubule regulator) [109–111], and *DTNBP1* (encoding the synaptic regulator dysbindin) [112, 113], improving recall at broader retrieval depth.

### Reasoning over SEMPER’s candidates improves target recovery

We further compare the LLM, SEMPER, and the LLM reordering on SEMPER predictions on the recovery of clinically pursued drug targets in Parkinson’s disease (Figure 5f), Alzheimer’s disease (Figure 5g), multiple sclerosis (Figure 5h), Huntington’s disease (Figure 5i), and amyotrophic lateral sclerosis (Figure 5j; Methods 6; Supplementary Note 2). We report recovery of held-out clinically pursued targets within the top 25 (Hits@25).

SEMPER recovers more clinically pursued targets than the LLM alone in all five diseases (Figure 5f-j). Across the 50 disease and cell type pairs (5 diseases × 10 cell types), SEMPER reaches a mean Hits@25 of 8.3 against 4.7 for the LLM (*p*-value = 1.7 × 10^−6^, Wilcoxon signed-rank test; Methods 6), showing that the LLM’s parametric knowledge recovers fewer held-out targets than SEMPER’s retrieval. Prompted only with the example targets, the LLM tends to nominate the well-characterized disease genes, which may not necessarily coincide with clinical or therapeutic relevance. For example, in amyotrophic lateral sclerosis, the LLM nominates well-established ALS-associated genes (*C9orf72* [114], *VCP* [115], *OPTN* [116], *SQSTM1* [117], *ATXN2* [118]) with no drug past early-phase trials, placing them outside our clinically pursued target set (Methods 1.5), resulting in the LLM recovering almost none of the held-out target set.

Allowing an LLM to reason over SEMPER’s top candidates improves recovery further over SEMPER alone, raising mean Hits@25 from 8.3 to 9.2 (*p*-value = 1.1 × 10^−2^), and exceeds the LLM alone by a wide margin (*p*-value = 5.9 × 10^−10^). The improvement from the LLM is largest in Huntington’s disease, where mean Hits@25 increases from 18.2 to 21.9 (*p*-value = 1.9 × 10^−3^). Parkinson’s disease (Hits@25 from 7.6 to 8.1), multiple sclerosis (Hits@25 from 5.6 to 5.9), and Alzheimer’s disease (Hits@25 from 5.5 to 5.6) show a smaller, non-significant upward shift in the Hits@25 distribution, and a slight decrease in amyotrophic lateral sclerosis.

The performance gain from LLM reordering varies across the five diseases (Figure 5f-j), and the rationales the LLM produces explain this variance. Because the LLM re-scores each candidate from its own parametric knowledge, we reason that reordering raises recovery when three conditions hold together. First, the pathology of the disease follows a mechanism that the literature describes in detail. Second, the clinically pursued targets belong to that mechanism and are among the candidates SEMPER retrieves. Third, the knowledge the LLM holds is specific enough to separate those targets from candidates that share the same mechanism without being targets.

Huntington’s disease satisfies all three conditions, as mitochondrial respiratory dysfunction is well established in Huntington’s [119–121], and its clinically pursued targets are the electron transport chain subunits. The LLM assigns high scores to Complex I subunits (e.g., MT-ND6, NDUFV1, NDUFS2, NDUFS8, and NDUFV2) [122] between 4.4 and 4.6 in all ten cell types. These scores are far above the 2.4 it assigns to non-targets, with the remaining high scores going to Complex I assembly factors [123–125] (NDUFAF6: 4.3; TMEM126B: 4.2) rather than unrelated proteins. Similarly, Parkinson’s disease is characterized by degeneration of the dopaminergic nigrostriatal system [6], and the LLM scores clinically pursued neurotransmitter receptor targets [126] (GRM5: 4.5; GRIA2: 4.3; GRIA1: 4.1) above generic synaptic proteins. However, competing non-target proteins in the same pathway (NTRK2: 4.1; VAMP1: 3.9) lower the LLM’s target assessment performance.

In Alzheimer’s disease and amyotrophic lateral sclerosis the first two conditions hold but the third fails, and the gain is smallest (Figure 5g-j). Alzheimer’s disease has a well-established amyloid pathology, but the mechanism the literature most often invokes, synaptic loss, applies to a large number of synaptic proteins [127, 128]. The LLM scores non-target amyloid- and synapse-associated proteins (EPHB2: 5.0; APBA2: 4.9; NTRK2: 4.7) at or above established glutamate receptor targets (GRIA2: 4.6; GRIA1: 4.4), failing to separate general disease-associated genes from genes/proteins pursued as drug targets. Amyotrophic lateral sclerosis is mechanistically heterogeneous [129], and the LLM’s predominant rationale invokes glutamate excitotoxicity, which applies equally to proteins that are and are not pursued as targets. The LLM scores well-known ALS-associated genes that are not clinically pursued targets [130–133] (*TUBA4A*: 5.0; *FIG4*: 4.9; *KIF5A*: 4.6; *UNC13A*: 4.4) above pursued targets such as *GRIN2D* (4.1), *PDE4D* (3.1), and *SCN9A* (1.3). The LLM’s biological rationales are consistent with known disease mechanisms, but many proteins share these mechanisms, so the rationales do not distinguish clinically pursued targets from functionally related proteins. The LLM ranks EPHB2 above the target GRIA2 in Alzheimer’s disease and TUBA4A above the target GRIN2D in ALS.

Multiple sclerosis shows only a modest improvement (5.6 to 5.9; Figure 5h) because SEMPER retrieves targets expressed in the CNS cell types it represents but misses many immunomodulatory targets that act in peripheral immune cells. Many clinically pursued multiple sclerosis targets act on immune mechanisms, including anti-CD20, S1P receptor, BTK, and BAFF/TNFSF13B agents, and on peripheral lymphocyte populations that are not among the brain cell types SEMPER represents [134–138]. These targets are not among the candidates SEMPER retrieves. For example, key target proteins such as *TNFSF13B, MS4A1, S1PR1*, and *BTK* are all absent, which limits recovery at the retrieval stage. The targets SEMPER does retrieve, and that the LLM scores highest, correspond to genes encoding neuronal ion channels (*SCN8A*: 4.9; *KCNA2*: 4.5; *KCNA1*: 4.4), while the high-scoring non-targets correspond to axonal and myelin genes (*NFASC*: 5.0; *NEFL*: 4.9; *CNTN1*: 4.9). The LLM thus refines SEMPER’s rankings rather than extending them, adding value where the targets are both retrievable from SEMPER and mechanistically separable from the proteins that resemble them.

### Gene annotations separate targets from functionally related genes

Scoring each gene on its own, from what the large language model already knows, leaves clinically pursued targets in Alzheimer’s disease and amyotrophic lateral sclerosis ranked below genes that share their mechanism (Figure 5g-j). We therefore give the large language model the annotated functions of each gene and ask it to compare genes against each other. We compare three approaches, the large language model reasoning from the example genes alone (Figure 6a), SEMPER’s cell type-specific retrieval (Figure 6b), and SEMPER’s retrieval reordered by the large language model using Gene Ontology annotations (Figure 6c). In the third approach, SEMPER’s embedding similarities set the initial rating of its top 100 genes, and a pairwise tournament then pairs genes of adjacent rank over successive rounds. For each pair, the large language model retrieves the Gene Ontology annotations of the two genes through a tool call [45, 46, 139] and decides which of the two is the more relevant target, and each gene’s rating is updated by its wins and losses (Figure 6c; Methods 6; Supplementary Note 3).

**Figure 6:**
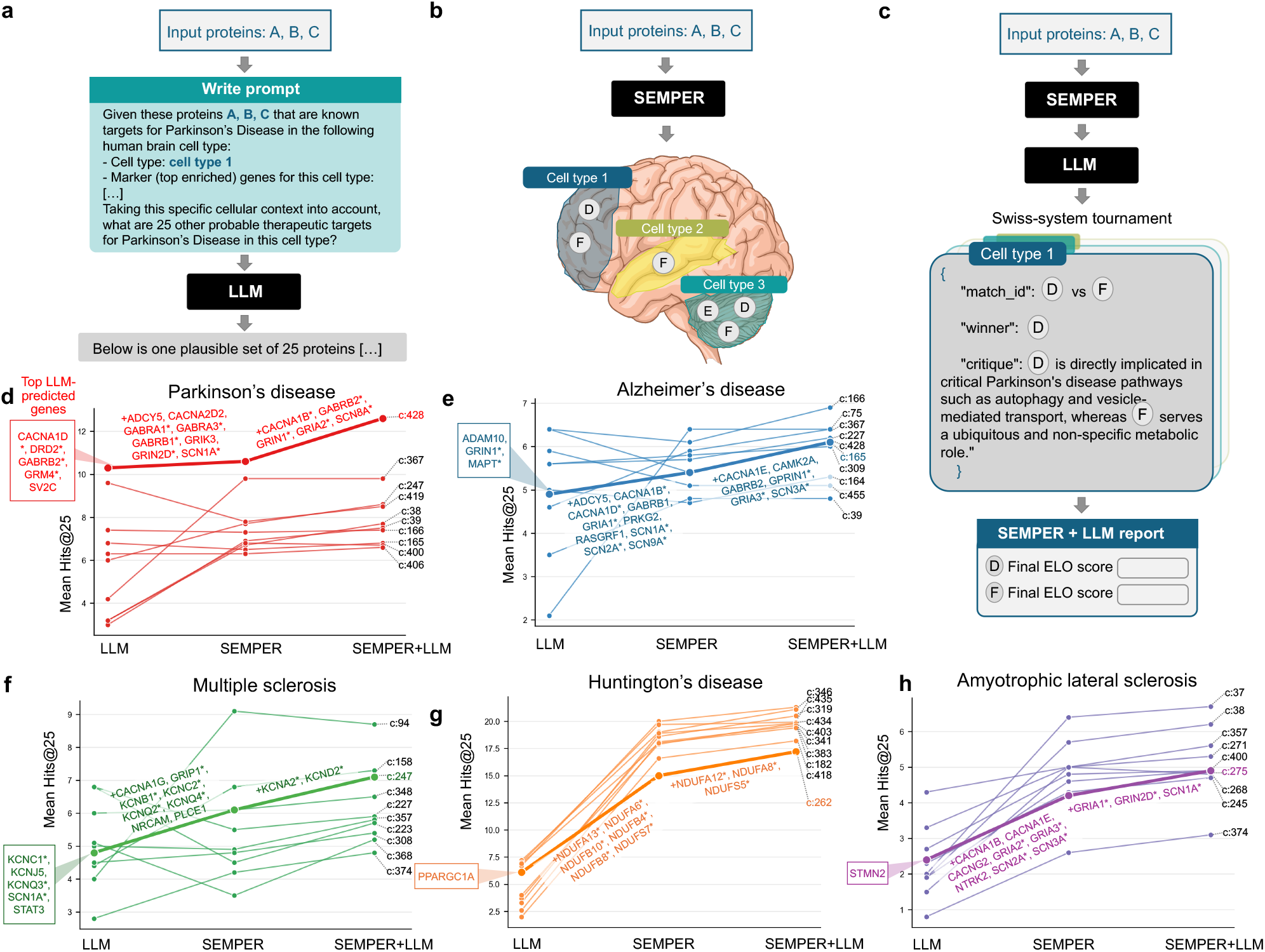
A tournament over SEMPER’s candidate genes recovers clinically pursued drug targets. **(a)** A reasoning large language model (GPT-5.6-sol) receives the seed targets and the cell type context (cell type identity and its top marker genes), and predicts additional candidate genes from its internal knowledge. **(b)** SEMPER retrieves cell type-specific candidate proteins from the seed genes. **(c)** A pairwise tournament that reorders SEMPER’s candidate genes. SEMPER’s similarities set the initial Elo ratings of its top 100 candidate genes. The genes then enter a multi-round Swiss-system tournament in which adjacently ranked genes are paired. For each pair, a large language model retrieves Gene Ontology annotations through a tool call, decides the winner from the annotated functions, and generates a biological rationale. Elo ratings are updated over successive rounds to produce the reordered gene list. **(d-h)** Slopegraphs of Hits@25 per cell type for the large language model alone, SEMPER, and SEMPER reordered by the tournament, across **(d)** Parkinson’s disease, **(e)** Alzheimer’s disease, **(f)** multiple sclerosis, **(g)** Huntington’s disease, and **(h)** amyotrophic lateral sclerosis, for the ten highest-recovery cell types from Figure 4. The bold slope marks the cell type whose recovery increases at each stage and improves most from SEMPER to the reordered ranking. For each highlighted cell type, the left box lists the genes the large language model placed first (rank-1) in at least one of the ten bootstrap runs. Genes marked with a ‘+’ prefix reached rank-1 under the reordered ranking and not under SEMPER alone. Asterisks (*) denote genes that are clinically pursued targets for the respective disease.

Reordering SEMPER’s candidate genes through the tournament improves recovery over the large language model alone and over SEMPER alone in all five diseases (Figure 6d-h). Across the 50 disease and cell type pairs, the tournament raises mean Hits@25 from 8.3 to 9.0 (*p*-value = 6.4 × 10^−9^, Wilcoxon signed-rank test). The increase holds in Parkinson’s disease, from 7.6 to 8.2 (*p*-value = 5.4 × 10^−3^; Figure 6d), Alzheimer’s disease, from 5.5 to 5.9 (*p*-value = 8.8 × 10^−3^; Figure 6e), multiple sclerosis, from 5.6 to 6.2 (*p*-value = 2.9 × 10^−3^; Figure 6f), Huntington’s disease, from 18.2 to 19.7 (*p*-value = 9.8 × 10^−4^; Figure 6g), and amyotrophic lateral sclerosis, from 4.8 to 5.1 (*p*-value = 2.9 × 10^−3^; Figure 6h). The reordered ranking also outperforms the large language model alone (9.0 versus 4.7; *p*-value = 2.2 × 10^−8^), including in Alzheimer’s disease and amyotrophic lateral sclerosis, where scoring each gene on its own left clinically pursued targets below genes of the same mechanism.

The annotations describe the specific functions of each gene, which a rationale stated at the level of the disease omits. In Alzheimer’s disease, where a synaptic rationale applies to many genes beyond the targets, the tournament places the glutamate receptor target genes *GRIA3* and *GRIA2* above *CAMK2A*, a calcium/calmodulin-dependent kinase, and *ADGRL3*, an adhesion receptor, on their annotated AMPA receptor activity and their localization to the postsynaptic density and dendritic spines. In amyotrophic lateral sclerosis, where an excitotoxicity rationale applies equally to the targets and to the genes of the same mechanism, the tournament places the sodium channel target genes *SCN1A* and *SCN2A* above the calcium channel genes *CACNA1B* and *CACNA1C* on their annotated roles in sodium-dependent action potentials. The same comparison places the NMDA receptor target gene *GRIN2D* above *CACNG2* on its annotation in NMDA-dependent calcium signaling. In each pairing, the comparison depends on a function annotated for the individual gene, so the reordered list ranks targets above the mechanistic neighbors that pointwise scoring placed ahead of them.

Target recovery for multiple sclerosis also improves (Figure 6f). Using the retrieved annotations, the tournament places the potassium channel target genes *KCNA2* and *KCND2* at rank 1 [140], and the comparison that places *KCNA2* there depends on its annotated localization to the juxtaparanodal region. The improvement is limited to the targets SEMPER represents. Disease-modifying treatments for multiple sclerosis are immunomodulatory and act on proteins expressed by peripheral lymphocytes, a population outside the CNS cell types of SEMPER. Target genes such as *TNFSF13B, MS4A1, S1PR1*, and *BTK* therefore fall outside SEMPER’s candidate lists, and the tournament reorders only the genes already on those lists. Across all five diseases, the tournament improves target recovery for the targets SEMPER retrieves and whose annotations describe a function that separates them from the genes of the same mechanism.

## Discussion

SEMPER is a multimodal representation learning model that integrates protein sequence with cell type-specific interaction context across 442 cell types of the adult human CNS. It learns a representation for each protein in every cell type expressing its gene, so representations vary across cellular environments while retaining the evolutionary information encoded in protein sequences. This integration improves therapeutic target prediction across protein language models from three ESM families, so the gain follows from the alignment and not from the choice of sequence model.

In SEMPER latent space, proteins are organized by cell type and biological function, and their representations predict CNS protein complexes under four definitions of cell type-specific assembly. SEMPER also predicts clinically pursued therapeutic targets across five neurodegenerative diseases and the genes essential to the survival of dopaminergic neurons. Across these analyses, representing proteins within their cellular contexts captures biological information beyond a single cell type-agnostic representation. Target recovery varies substantially across cell types, so the representations distinguish the cellular contexts in which the targets of a disease are most readily recovered.

The information in SEMPER and the knowledge a large language model holds are complementary, and each improves the other. Large language models depend on cell types described in the published literature, which limits the range of cell types within reach of their reasoning. SEMPER characterizes cellular context from experimental measurements instead, so a large language model reasoning over SEMPER’s candidate genes reaches cell types with sparse literature coverage. In the other direction, published knowledge improves SEMPER’s assessment of clinically relevant genes when it describes mechanisms that separate the targets from related proteins. That condition holds in Huntington’s disease and Parkinson’s disease, where established mechanisms such as mitochondrial Complex I dysfunction and *α*-synuclein biology separate the targets from the rest of SEMPER’s candidate genes, and these are the diseases with the largest improvement. The condition fails for broader mechanisms such as synaptic loss in Alzheimer’s disease and glutamate excitotoxicity in amyotrophic lateral sclerosis, which apply to many genes and are less informative about which of those genes are clinically pursued targets. Protein functional annotations restore the distinction by making the reasoning specific to each gene, which improves target recovery across all five diseases.

SEMPER’s cellular resolution follows the coverage of the single-cell atlas, including the granularity of the human-annotated cell types. The 442 cell types are the clusters that annotation defines, so a population divided into two clusters in one atlas and one cluster in another receives correspondingly different representations. Because the protein representations are learned from CNS interaction networks constructed from that atlas, proteins in sparsely sampled cell types, or proteins with poorly characterized interactions, have less reliable representations, since the interaction neighborhood that distinguishes a protein in one cell type from the same protein in another is estimated from fewer measurements. SEMPER represents proteins only within the cellular contexts present in these networks. Multiple sclerosis shows the consequence, since many disease-modifying targets act on peripheral immune cell populations outside SEMPER’s coverage, which places those targets outside its candidate genes in the zero-shot retrieval analysis and outside the set of genes available to subsequent reasoning. Extending the networks to peripheral and immune cell types would provide representations for proteins acting in those contexts and make their disease biology available to the same analyses.

A second extension concerns the order of the two models. At present a large language model reasons only over the genes SEMPER retrieves, so the final ranking inherits SEMPER’s coverage, and a target the literature describes well is unavailable to the reasoning step whenever SEMPER does not retrieve it. Letting SEMPER and a large language model propose candidate genes in parallel would produce two sets of genes, each covering genes outside the other.

The zero-shot retrieval benchmark evaluates recovery of targets that already carry clinical evidence and ChEMBL mechanism annotations. The benchmark therefore measures whether the representations place known targets near one another, and it leaves open the prioritization of therapeutic targets that are currently unrecognized, since a benchmark built from pursued targets provides no measurement of the genes that have yet to be pursued. Clinical pursuit is also distinct from therapeutic efficacy, particularly for targets still under clinical evaluation. Prospective evaluation of newly prioritized genes will determine whether SEMPER extends beyond the recovery of previously pursued targets. Embedding proximity is associative, so SEMPER’s predictions are hypotheses whose biological effects require experimental testing.

SEMPER integrates evolutionary information from protein sequences with cell type-specific interactions to represent protein biology across CNS contexts. SEMPER supports analyses of molecular complexes, candidate therapeutic targets, and essential genes across CNS cell types and diseases. Across these analyses, SEMPER represents cell type dependent biological relationships more accurately than models that learn from protein sequences alone, from cellular context alone, or from the published literature alone. Resolving proteins by the cell types in which they act, while keeping them grounded in their sequences, changes which aspects of protein function are open to prediction, and we expect this to support the study of neurodegenerative diseases in the cell populations that are selectively vulnerable to them.

## Acknowledgements

We thank Richard Zhu for helpful discussions on parts of this manuscript.

## Data availability

Benchmark datasets are available at https://github.com/mims-harvard/SEMPER/tree/main/data/benchmarks. They include the 547 CNS-relevant CORUM protein complexes (Supplementary Table 4) and, for each of the four complex definitions, the complexes present in each cell type together with the size-matched decoys used for evaluation. They also include the clinically pursued therapeutic targets for Alzheimer’s disease, Parkinson’s disease, multiple sclerosis, Huntington’s disease and amyotrophic lateral sclerosis, with the clinical phase, associated drugs, pharmacological action and therapeutic intent reported for each target (Supplementary Table 6), and the 693 genes essential to the survival of human iPSC-derived midbrain dopaminergic neurons identified in the genome-wide CRISPR screen of Nazeen et al. [89]. The cell type-agnostic interactome TransposeNet 2 is available at Figshare [141]. The single-nucleus transcriptomic atlas of the adult human brain [38], the brain tissue protein co-abundance atlas [52] and the Parkinson’s disease clinical trial dataset [82] are available from their original publications. CORUM [142], the Human Protein Atlas [56], Gene Ontology [45], the Open Targets Platform [143], ChEMBL, DrugBank, UniProtKB [144] and Ensembl [145] are available from their respective portals. The 442 cell type-specific PPI networks, the metagraph and the SEMPER protein representations are available from the corresponding author upon reasonable request. The project website is at https://zitniklab.hms.harvard.edu/SEMPER.

## Code availability

SEMPER is available at https://github.com/mims-harvard/SEMPER under MIT license.

## Author contributions

A.D., M.M.L., and M.Z. developed the protein representation learning model. A.D. implemented the model and executed the analyses. A.D., M.M.L., and M.Z. designed the evaluations of the networks and the model. M.M.L., K.D., S.N., and A.N. constructed the cell type specific protein-protein interaction networks. Y.H. contributed to the protein language model analyses. S.G. contributed to the agent and tool-use components. D.R., J.A., A.B.H.H., and L.S. conducted and analyzed the genome-wide CRISPR/Cas9 screen in midbrain dopaminergic neurons. Y.L. contributed to the network construction. G.P. and M.M.L. designed the statistical analyses. V.K. provided feedback and discussion on the analyses related to neurodegenerative diseases. A.D., M.M.L., and M.Z. wrote the manuscript with input from all co-authors. M.Z. led the overall research study.

## Online Methods

The Online Methods describe the construction of training and evaluation datasets (Methods 1); the SEMPER architecture and multimodal contrastive alignment (Methods 2); the benchmarking of SEMPER’s latent space and cell type-specific protein complex prediction (Methods 3); the zero-shot retrieval of candidate drug targets for neurodegenerative diseases (Methods 4); the assessment of essential genes in dopaminergic neurons (Methods 5); and the LLM and statistical analyses used (Methods 6).

## 1 Datasets

### 1.1 Construction of multi-scale protein-protein interaction networks

SEMPER’s multi-scale networks consist of 442 cell type-specific PPI networks and one unifying cell-tissue metagraph. The PPI networks encode physical protein-protein interactions restricted to the proteins active in each cell type. The metagraph encodes the anatomical and taxonomic hierarchy that relates cell types, superclusters, brain structures, and brain regions.

#### Cell type-specific protein-protein interaction networks

We construct a cell type-specific protein-protein interaction (PPI) network for each of the 442 cell types (clusters) annotated in a single-nucleus transcriptomic atlas of the adult human brain (24,501 genes assayed across 2,399,168 neurons and 970,051 non-neuronal cells from 3 donors) [38, 146], following the procedure of PINNACLE [20]. For each cell type, we first select *k* = 4,000 differentially expressed genes (DEGs; *P* ≤ 0.01) [146] that capture both the cell type’s broad and fine-grained identity. Half of the genes (*k*/2) are the top DEGs distinguishing the cell type’s supercluster from all other superclusters, with each cell type belonging to one of 30 superclusters. The other half of the genes are the top DEGs distinguishing the cell type from the other cell types within its supercluster. We then map these genes to their proteins, extract the subgraph induced by these proteins from a cell type-agnostic PPI network [39, 141] (Methods 1.2), and keep the largest connected component as the cell type-specific PPI network.

#### Cell type and tissue relationships in the metagraph

The metagraph represents the organization of a root node (i.e., whole brain), 14 brain regions (i.e., amygdaloid complex, basal forebrain, basal nuclei, cerebellum, cerebral cortex, claustrum, extended amygdala, hippocampus, hypothalamus, midbrain, myelencephalon, pons, spinal cord, thalamus), 112 brain structures, 30 superclusters, and 442 cell types represented in the single-cell transcriptomic atlas of the adult human brain [38, 146]. 2,316 edges capture the hierarchical relationships among these 599 nodes.

#### Data statistics of final multi-scale networks

The 442 cell type-specific PPI networks contain an average of 2,843 ± 134 proteins, ranging from 2,385 to 3,146. Summed across cell types, they contain 1,256,780 protein nodes, which correspond to 15,487 unique proteins appearing in at least one cell type. Each of these 15,487 proteins therefore has one amino acid sequence and, on average, 81 cell type-specific network contexts. The metagraph contains 599 nodes and 2,316 edges.

#### Evaluation of cell type-specific PPI networks

To assess whether the resulting cell type-specific PPI networks capture biologically meaningful brain-specific protein associations, we compare network edges against an independent brain tissue protein co-abundance atlas derived by Laman Trip et al. from 7,811 human proteomic samples [52]. Within each cell type-specific PPI network, we compare connected pairs, which are pairs of proteins linked by an edge, with unconnected pairs, which are pairs of proteins that are both present in the network but not linked, using only pairs scored in the atlas. Connected pairs have higher brain co-abundance than unconnected pairs in all 442 cell types (median 0.342 versus 0.288; one-sided Wilcoxon signed-rank test on paired per-cell type medians, *p*-value= 1.9 × 10^−74^; Supplementary Figure 2). The cell type-specific PPI networks therefore agree with independently measured protein co-abundance.

### 1.2 Cell type-agnostic PPI network

The cell type-agnostic PPI network is a humanized, genome-scale PPI network derived from TransposeNet 2 [39, 141]. TransposeNet 2 is a cross-species homology-augmented network of human PPIs. Its human component merges STRING v12.0 [147] (restricted to only experimental and database evidence channels) with three resources of high-quality binary protein interaction maps: the Human Reference Interactome (HuRI) [148], Lit-BM-13, and HI-II-14 (the latter two from Rolland et al. [149]). An interactome of baker’s yeast (*Saccharomyces cerevisiae*) is assembled in the same manner from STRING v12.0 and ABBI-21 [150]. Similarly to the original TransposeNet [39] framework, TransposeNet 2 augments the human network first by mapping interacting yeast proteins to their human homologs using a hierarchy of evidence: sequence similarity (BLAST [151]), 3D structural similarity (Foldseek [152]), profile-HMM similarity (HHpred [153]), curated orthology (DIOPT [154]), and topological similarity (Mashup [39, 155]). Then, every yeast interaction whose two partners both carried homologs was transferred to the corresponding human protein pairs. This augmentation either introduces a new edge or strengthens the evidence of an existing one in the original human PPI network. Edges with a confidence score ≥ 0.5 are retained, and the largest connected component of the interactome is extracted, resulting in an interactome of 17,032 proteins and 444,330 interactions, of which 161,277 (36.3%) are homology-transposed from yeast. The constituent data sources and the quality of edges are systematically evaluated as part of the construction and validation of TransposeNet 2 [141]. Full details of the network construction and validation will be provided in the forthcoming TransposeNet 2 publication.

### 1.3 Tissue-specific protein co-abundance atlas

To assess whether SEMPER’s representations reflect experimentally measured, brain-specific protein co-abundance, we leverage the tissue-resolved protein co-abundance atlas of Laman Trip et al. [52] as a source of orthogonal validation. The atlas reports protein co-abundance across 11 human tissues (brain, ovary, lung, liver, pancreas, blood, breast, throat, colon, stomach, and kidney), computed from mass spectrometry proteomics of 7,811 tumor and tumor-adjacent human biopsy samples. Within each tissue, the atlas assigns every pair of co-measured proteins a co-abundance association score, defined as the average probability that the two proteins are co-abundant in that tissue ranging from 0 (no evidence of co-abundance) to 1 (strong, consistent co-abundance).

Following Laman Trip et al. [52], we consider a protein pair to be specific to a given tissue when their co-abundance score exceeds the 95th percentile of that tissue’s score distribution (0.68 ± 0.01 across tissues) while remaining below 0.5 in all other tissues. After we map the atlas’s gene identifiers to SEMPER’s proteins, SEMPER and the brain co-abundance atlas share 10,621 proteins (68.6% of SEMPER’s 15,487 proteins; 5,885–12,053 proteins for the other ten tissues), and we perform all co-abundance analyses on protein pairs within this shared set.

We use two thresholds on this atlas. The 95th-percentile rule above defines tissue-specific pairs for the latent space analysis (Methods 3.1.4). A flat score of 0.5, below which Laman Trip et al. consider a pair not co-abundant, defines the protein co-abundance complex criterion (Methods 1.4).

### 1.4 Curation of CNS-relevant protein complexes

We retrieve human protein complexes from the Comprehensive Resource of Mammalian Protein Complexes (CORUM) database [142]. Because CORUM is not cell type-resolved, we first restrict the dataset to complexes with documented central nervous system relevance. For all canonical human entries, we provide the complex name, source publications, curator comments, and Gene Ontology annotations supplied by CORUM to the o3-mini reasoning model [156], and score each complex on a scale of 1 to 5 for evidence of relevance to CNS biology, neurodevelopment, and neurodegenerative disease. The full prompt and rubric appear in Supplementary Note 1. We retain complexes that score ≥ 4, contain at least three distinct subunits, and have every subunit present in the PPI network, yielding a final benchmark of *n* = 547 CNS-relevant complexes (refer to Supplementary Table 4 for a full list of CNS-relevant complexes).

#### 1.4.1 Defining cell type-specific complex benchmarks

A protein complex can assemble only where all of its subunits are present in the same cell and the same subcellular location [53, 54]. A CORUM complex therefore may not assemble in every cell type in which its subunits are expressed, because co-expression is necessary but not sufficient for assembly. We define four criteria for defining a complex as present in a given cell type. The base criterion, transcriptional co-expression, requires all subunits to be expressed in that cell type. Three strict criteria, structural connectivity, protein co-abundance, and spatial colocalization, each add one further line of evidence to the base criterion. Because a complex that meets one strict criterion need not meet another, we treat each definition as a separate benchmark.

The base criterion, transcriptional co-expression, requires every subunit of a CORUM complex to be present in the cell type’s PPI network. The second definition, structural connectivity, additionally requires the complex subunits to be linked to one another by physical interactions in the cell type, such that the subgraph of the cell type’s PPI network induced by the subunits is a connected component. The third definition, protein co-abundance, requires every pair of subunits to have a brain co-abundance score of at least 0.5 in the atlas of Laman Trip et al. [52] (Methods 1.3), and excludes complexes with any subunit pair missing from the atlas. The fourth definition, spatial colocalization, requires all subunits to share at least one subcellular location annotated in the Human Protein Atlas (HPA) [56], and excludes complexes with any subunit lacking an HPA annotation.

#### 1.4.2 Negative sampling strategy

To turn each definition into a binary classification task, we generate decoy complexes at a 1:10 positive-to-negative ratio for every cell type and definition. Each decoy matches the subunit count *k* of its corresponding positive complex, so that no model can separate positives from negatives by complex size alone, and contains only proteins present in that cell type’s network. All decoys are de-duplicated against the entire CORUM database to ensure no randomly sampled decoy is equivalent to a canonical biological complex. For the spatial colocalization definition, we sample each decoy of size *k* only from proteins that share the same HPA subcellular annotation, so that decoys match positives in subcellular location.

### 1.5 Curation of candidate therapeutic target genes

We curate clinically pursued targets for five neurodegenerative diseases: Alzheimer’s disease (AD), Parkinson’s disease (PD), multiple sclerosis (MS), Huntington’s disease (HD), and amyotrophic lateral sclerosis (ALS). We utilize the Open Targets Platform [143], a comprehensive biomedical informatics database that integrates genetic, genomic, and drug data. For each disease (e.g., Alzheimer’s disease), we identify clinical trials for the condition and its descendants in the disease ontology (e.g., early-onset Alzheimer’s disease). Among the genes investigated in these trials, we filter for protein-coding genes that have been evaluated as candidate targets in completed phase 2 trials or in phase 3–4 trials of any status. We report the number of targets by trial phase and classify their associated drugs by therapeutic intent as symptomatic or disease-modifying (Fig. 4a). We map HGNC symbols [157] to Ensembl identifiers.

The curation yields 130 clinically pursued targets for Alzheimer’s disease, 97 for Parkinson’s disease, 166 for multiple sclerosis, 86 for Huntington’s disease, and 98 for ALS (Supplementary Table 6). Because retrieval runs within each cell type-specific network (Methods 4), only the targets present in a given network are recoverable there. Among cell types whose networks contain more than 40 targets, the number of targets present averages 45.6 for Alzheimer’s disease (191 cell types; range 41 to 61), 44.0 for Parkinson’s disease (98 cell types; range 41 to 56), 49.6 for multiple sclerosis (324 cell types; range 41 to 69), 50.8 for Huntington’s disease (143 cell types; range 41 to 61), and 42.4 for ALS (17 cell types; range 41 to 47). Refer to Supplementary Table 6 for the full target list per disease.

#### Validation of curated clinically pursued PD targets

To assess our pipeline for curating clinically pursued targets, we compare our clinically pursued targets for Parkinson’s disease against an independent, manually curated dataset of 444 Phase 1-3 Parkinson’s disease drug trials registered on ClinicalTrials.gov between 2015 and 2024 [82]. To match our definition of clinically pursued targets, we restrict the comparison to the 141 trials in Buff et al. that have completed Phase 2 or are in Phase 3 of any status, following Buff et al. in treating Phase 2/3 trials as Phase 2. Because Buff et al. reports agents and mechanism classes but not target genes, we map each trial’s agents to protein targets using ChEMBL mechanism-of-action annotations, the mechanism subcategories that Buff et al. assign (e.g., dopamine agonist, COMT inhibitor), and for the three trials whose agents we could not link to a ChEMBL mechanism, their published mechanism. Excluding 22 trials of therapies without a defined protein target (e.g., cell, gene, and microbiome therapies) and two trials with undisclosed mechanisms, 105 of the remaining 117 trials (90%) test an agent that acts on at least one of our targets, including 94 of 99 symptomatic trials (95%) and 11 of 18 disease-modifying trials (61%). Of the remaining 12 trials, 10 test agents that lack ChEMBL mechanism-of-action annotations and 2 test agents for which Parkinson’s disease (or its descendants in the disease ontology) is not listed as an indication according to Open Targets. Together, the 105 overlapping trials involve 62 of our 97 targets (64%). Because Buff et al. excludes Phase 4 trials, targets of approved drugs appear only through reformulations, new agents, or repurposing; these trials’ agents act on 24 out of 34 (71%) targets from our curated Phase 4 trials.

## 2 Overview of the SEMPER Model

SEMPER maps cell type-specific protein-protein interactions and cell type-agnostic protein sequences into one shared latent space. The model uses two pretrained modality encoders. A multi-scale graph transformer encodes the 442 cell type-specific PPI networks together with the metagraph that nests cell types within superclusters, brain structures, and brain regions. A protein language model encodes each protein’s amino acid sequence. SEMPER freezes both encoders during multimodal alignment and trains only two projection heads, which map each modality into a shared 512-dimensional space under a symmetric InfoNCE contrastive loss [41, 42]. SEMPER forms a protein’s final representation in a given cell type by combining its sequence projection with its interaction projection for that cell type in that shared space.

### 2.1 Modality encoders

SEMPER uses a dual-encoder architecture analogous to CLIP [158], with one branch per modality. The interaction branch is a multi-scale graph transformer that processes the 442 cell type-specific PPI networks together with the metagraph and returns one representation for each protein in every cell type where that protein is expressed. The sequence branch is a pretrained protein language model that reads a protein’s amino acid sequence and returns a single, cell type-agnostic representation per protein. Because the interaction branch yields many representations per protein and the sequence branch yields one, this asymmetry shapes how SEMPER pairs the two modalities during alignment.

#### 2.1.1 The multi-scale network encoder

SEMPER uses a multi-scale graph transformer encoder to learn cell type-specific PPI representations of proteins. The encoder jointly models (i) the 442 cell type-specific PPI networks and (ii) the metagraph that relates cell types and tissues.

#### Problem formulation

Let *P* denote the set of proteins in the unfiltered cell type-agnostic interactome, *C* the set of 442 cell types, and *T* the set of tissue nodes of the metagraph (Methods 1.1). For each cell type *c* ∈ *C*, we define a cell type-specific PPI graph *G*_*c*_ = (*V*_*c*_, *E*_*c*_), whose nodes *V*_*c*_ ⊆ *P* are the proteins in the network of cell type *c* and whose edges *E*_*c*_ ⊆ *V*_*c*_ × *V*_*c*_ are the interactions among them.

We also define a single directed metagraph *G*_*M*_ = (*V*_*M*_, *E*_*M*_), where the subscript *M* denotes the metagraph, with node set *V*_*M*_ = *C* ∪ *T* . Metagraph edges represent *R* = 4 relation types between nodes, namely cell type to cell type, cell type to tissue, tissue to cell type, and tissue to tissue, which we index by *r* ∈ {1, … , *R*}. We split the metagraph edges by relation type into subsets *E*_*r*_ , so that the encoder processes each kind of connection separately. In contrast, each PPI graph *G*_*c*_ contains a single relation type, protein to protein (*R* = 1).

#### Node features and hidden representations

We initialize a cell type-agnostic protein feature matrix 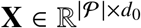 , with one row for each of the |*P*| = 19,097 proteins and initial feature dimension *d*_0_ = 512, using entries drawn i.i.d. from *N* (0, 1), and keep **X** fixed during training. For each cell type *c*, the feature matrix 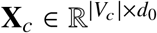 contains the rows of **X** for the proteins in *V*_*c*_, so each protein starts with the same features in every cell type. We pad the features to a dimension divisible by the number of attention heads. Within each encoder block *l* ∈ {0, 1}, we denote by 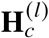 and 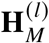 the input representations of the nodes in *V*_*c*_ and *V*_*M*_, respectively, by **Ĥ** ^(*l*)^ and 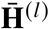 the representations after the first and second rounds of intra-graph message passing, and by 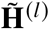 the output of the block, with 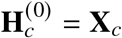. Metagraph nodes have no input features. Instead, the first encoder block constructs their representations from the protein representations, as described below. The output of the second block, 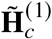, gives the final protein representations of the network encoder.

#### Overview of the hybrid graph transformer

SEMPER stacks two encoder blocks (*l* ∈ {0, 1}) built on the General, Powerful, Scalable (GPS) graph transformer [40]. Each block has separate weights for every cell type-specific PPI graph *G*_*c*_ and one shared set of weights for the metagraph *G*_*M*_. A block proceeds in four steps. The encoder first updates the proteins in each *G*_*c*_ with a GPS layer, which combines local message passing along PPI edges using GATv2 [159] with global multi-head attention across all proteins in that cell type. It then pools each cell type’s proteins into the corresponding cell type node of *G*_*M*_ as an attention-weighted sum of their representations, with weights *γ*_*c*_ (*p*) computed from each protein’s representation. Next, it propagates cell type information to the tissue nodes and updates all metagraph nodes with a GPS layer applied separately to each of the four relation types, combining the results by semantic attention. Finally, it updates the proteins in each *G*_*c*_ with a second GPS layer and adds to each protein its cell type node’s representation, scaled by the same weight *γ*_*c*_ (*p*) used for pooling. Between blocks, LayerNorm [160], LeakyReLU [161], BatchNorm [162], and dropout [163], applied in that order, transform 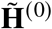into **H**^(1)^. The following sections describe each step in detail.

#### Metagraph representations

The two blocks construct the metagraph representations differently. In the first block, the encoder builds metagraph representations on every forward pass. It sets each cell type node to the attention-weighted sum of its proteins’ representations after the first round of message passing, using the weights *γ*_*c*_ (*p*) defined below, and initializes each tissue node with noise drawn from *N* (0, 1). In the second block, the encoder applies a metagraph GPS layer before pooling, and adds the newly pooled protein representations to the cell type nodes from the first block rather than reinitializing them. In both blocks, the encoder then propagates cell type information to the tissue nodes before metagraph message passing, replacing each tissue node’s representation with the mean of its neighbors in the undirected metagraph for 100 iterations,

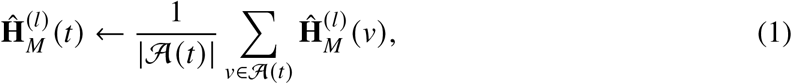

where *A*(*t*) denotes the set of neighbors of tissue node *t* in the undirected metagraph.

#### Intra-graph updates via semantic attention

The encoder applies the same update to the PPI graphs and the metagraph, so we write **H**^(*l*)^ for the node representations of any one graph at block *l*, standing for either 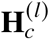 or 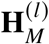. The update first processes each relation type separately. For each relation type *r*, a GPS layer combines local message passing along the edges in *E*_*r*_ using GATv2 [159] with global multi-head attention across all nodes of the graph [164], followed by a LeakyReLU activation and a linear projection, which together produce a relation-specific representation 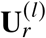

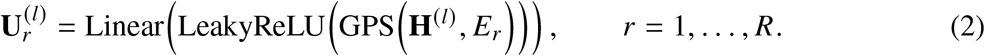

A semantic attention mechanism then combines the *R* relation-specific representations of each node *v* into a single representation

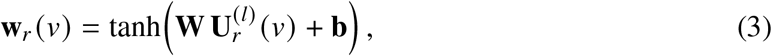

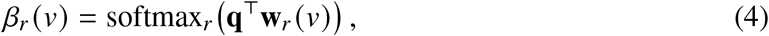

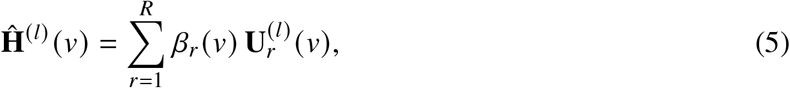

where **W, b**, and **q** are learnable parameters and *β*_*r*_ (*v*) weights the contribution of relation *r* to node *v*. The encoder applies this update to all proteins in each cell type, yielding 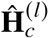 , and to all metagraph nodes, yielding 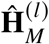. For the PPI graphs, which have a single relation type, *β*_1_(*v*) = 1 and the update reduces to the GPS layer, activation, and projection. The second round of protein message passing in each block applies the same update with separate weights, yielding 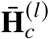.

#### Bridge between protein and cell type representations

The encoder links the protein and metagraph scales with a single set of protein-to-cell type attention weights, which it uses in both directions. After the first round of protein updates, it computes, for each protein *p* in cell type *c*,

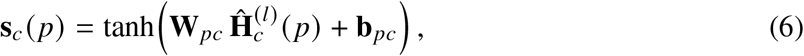

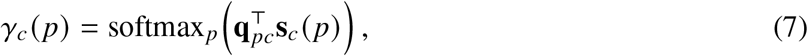

where **W**_*pc*_, **b**_*pc*_, and **q**_*pc*_ are learnable parameters and the softmax runs over the proteins of cell type *c* in the current mini-batch. The encoder first uses these weights to pool proteins into their cell type node,

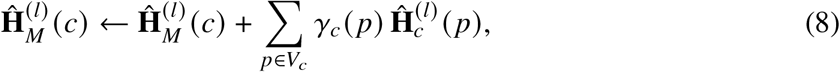

where 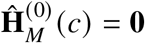 before pooling in the first block, so that pooling initializes each cell type node, and the second block adds to the existing representation. After tissue propagation and metagraph message passing produce the metagraph output 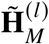 , the encoder applies the second round of protein updates, yielding 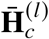 . It then uses the same attention weights in the reverse direction to add each cell type’s representation to its proteins,

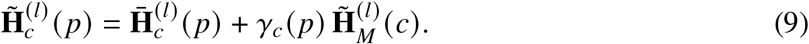

Reusing *γ*_*c*_ (*p*) in both directions ties how strongly a protein contributes to its cell type’s representation to how strongly it receives cell type context.

#### 2.1.2 Pretraining the multi-scale network encoder

Before multimodal contrastive alignment, we pretrain the network encoder with two self-supervised objectives, link prediction at both the PPI and metagraph scales and a protein-level center loss that encodes cell type identity. We write 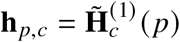 for the encoder’s final representation of protein *p* in cell type *c*, and 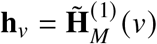 for that of metagraph node *v*, both of dimension *d*.

##### Data split

To evaluate the model’s performance without data leakage, we randomly split the protein-protein edges of each cell type-specific PPI graph G_*c*_ into training (80%), validation (10%), and test (10%) sets. Because the metagraph contains few edges and each is needed to preserve its hierarchy, the full metagraph remains visible during training and evaluation.

##### Mini-batch training

Processing the 1.25 million protein nodes within the multi-scale graph simultaneously is computationally intractable, so we train the encoder on mini-batches sampled with the GraphSAINT [165] edge sampler. At each training step, we sample from every cell type-specific PPI graph a subgraph made up of a subset of its edges and their proteins, and process the metagraph in full. We reset the sampled proteins’ features to their initial values **X**_*c*_ at the start of the batch, so that features that have undergone different amounts of message passing do not mix across batches. The encoder then runs its intra-graph and bridge steps, computes the link prediction and center losses, and updates its parameters by backpropagation. Node states reset with each batch, whereas the learned weights carry over.

##### Protein-level objectives

At the protein level, we train the encoder with link prediction within each cell type-specific PPI graph and with a center loss that aligns each protein’s representation with its cell type identity. For link prediction, we score each candidate edge between proteins *p* and *p*^′^ in cell type *c* with the dot product of their representations,

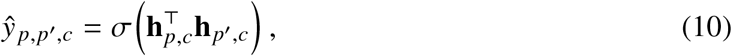

and minimize the binary cross-entropy against held-out true edges and an equal number of randomly sampled non-edges,

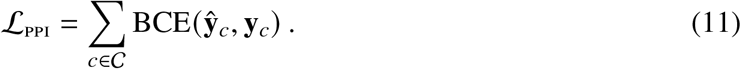

For cell type identity, we apply a center loss [166] that pulls each protein’s representation toward the metagraph representation of its cell type. Unlike the original formulation, the centers are not free parameters but the cell type representations **h**_*c*_ produced by the metagraph branch, which ties the protein and metagraph scales together,

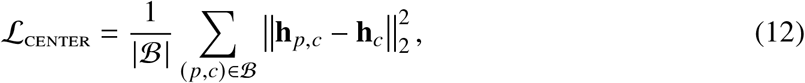

where B is the set of training protein-cell type pairs in the current mini-batch.

##### Metagraph-level objective

At the metagraph level, we train the encoder with link prediction over four relation types: cell-cell, cell-tissue, tissue-cell, and tissue-tissue. We score each candidate edge using metagraph nodes *v* and *v*^′^ of relation type *r* with a relation-specific elementwise gate **g**_*r*_ ∈ ℝ

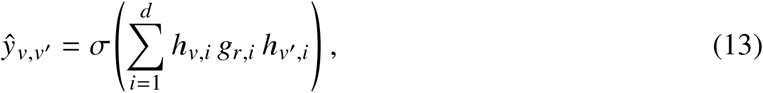

where the gates **G** ∈ ℝ^*R*×*d*^ are initialized with Xavier initialization [167] and kept fixed during training. The metagraph loss is the binary cross-entropy ℒ_mg_ = BCE(**ŷ**_*M*_, **y**_*M*_).

##### Overall objective

We combine the protein-level and metagraph-level losses as

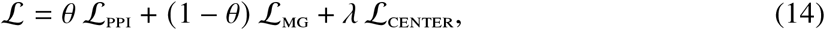

with *θ* = 0.1 and *λ* = 0.01, which jointly optimizes connectivity at both scales while enforcing cell type identity at the protein level.

#### 2.1.3 The protein sequence encoder

To represent each protein’s amino acid sequence, we use the ESM Cambrian (ESM-C) protein language model [12, 13]. ESM-C is trained on large sequence databases, including UniRef [168], MGnify [169], and the Joint Genome Institute [170], and learns a deep generative understanding of protein structure and function solely from evolutionary sequence patterns. Unlike the network encoder, which produces a separate representation for each cell type, ESM-C produces one representation per protein. We use the pretrained ESM-C as a frozen feature extractor and do not fine-tune on our CNS biological networks. We run inference exclusively on the canonical amino acid sequences corresponding to the unique proteins present in our multi-scale networks.

##### Data acquisition

We retrieve a canonical amino acid sequence for each protein in the cell type-agnostic interactome. Because the interactome mixes Ensembl protein (ENSP) [145] identifiers and UniProt [144] entry names, we handle the two separately. We retrieve sequences for ENSP identifiers through the Ensembl REST API [145], mapping deprecated identifiers to their latest stable successors through the Ensembl Archive. We map UniProt entry names to their primary accessions through the UniProtKB [171] search endpoint and download the corresponding sequences. We remove residual stop codon characters (`*`) before tokenization. This yields sequences for 18,680 proteins, covering every protein in the 442 cell type-specific networks.

##### Model inference

We embed each sequence with the `esmc_600m` model (600 million parameters) [13], whose hidden states have dimension 1,152. We compute the sequence representation **e**_*p*_ of protein *p* by mean-pooling the final hidden states **e**_*p, j*_ over its residues,

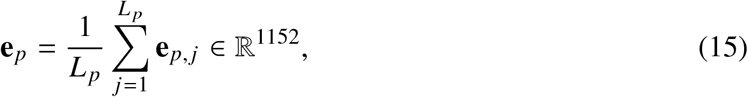

where *L* _*p*_ is the length of the sequence, excluding the beginning- and end-of-sequence tokens. Each protein therefore has exactly one sequence representation, shared across all cell types, which serves as the cell type-agnostic input to multimodal alignment.

### 2.2 Multimodal contrastive alignment

SEMPER aligns the sequence and network representations in a shared 512-dimensional space. We keep the pretrained network encoder and ESM-C frozen and train only two projection heads with a symmetric InfoNCE loss [41, 158], which pulls together the sequence and network representations of the same protein and pushes apart those of different proteins.

#### 2.2.1 Training pairs

For each cell type *c* and protein *p* ∈ *V*_*c*_, the pretrained network encoder provides a cell type-specific representation **h**_*p,c*_ ∈ ℝ^64^ (Methods 2.1.2), and ESM-C provides a single sequence representation **e**_*p*_ ∈ ℝ^1152^ (Methods 2.1.3). Because each protein has one sequence representation but a network representation in every cell type whose network contains it, we pair **e**_*p*_ with each of its network representations. The training set contains one pair (**e**_*p*_, **h**_*p,c*_) for every protein *p* with a retrieved sequence and every cell type *c* with *p* ∈ *V*_*c*_.

#### 2.2.2 Projection heads

Two multilayer perceptrons (MLPs) [172] with ReLU activations [173] map each modality into the shared space. The sequence head *f*_seq_ maps ℝ^1152^ to ℝ^512^ through hidden layers of 1,024 and 512 units, and the network head *f*_net_ maps ℝ^64^ to ℝ^512^ through two hidden layers of 512 units. Both heads end in a linear layer, and we ℓ_2_-normalize their outputs,

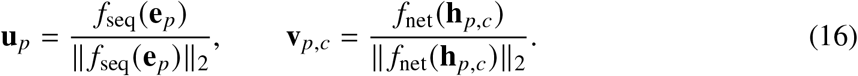

#### 2.2.3 Symmetric InfoNCE loss

We train the projection heads on mini-batches ℳ of *M* = 64 pairs. For pairs *j* and *j*^′^ in a mini-batch, we compute the scaled cosine similarity 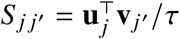 between the sequence projection of pair *j* and the network projection of pair *j*^′^, with a fixed temperature *τ* = 0.07. In each direction, the loss treats the matching pair as the correct class among the *M* candidates,

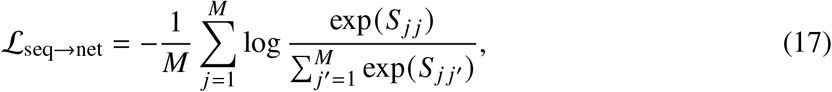

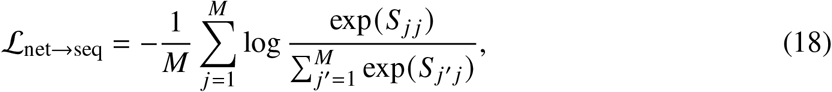

and the alignment loss is their average, 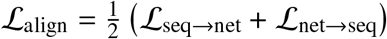. We train for 200 epochs with the Adam optimizer [174] at a learning rate of 10^−4^.

#### 2.2.4 Cell type-resolved representations

After training, we project every protein in every cell type. For the 243 of 15,487 network proteins without a retrieved sequence, which account for 16,872 of the 1,256,780 protein-cell type pairs (1.3%), we replace **u**_*p*_ with the normalized mean of the sequence projections of all proteins with retrieved sequences. Because the sequence projection **u**_*p*_ is the same in every cell type, we form the representation of protein *p* in cell type *c* by combining it with the cell type-specific network projection,

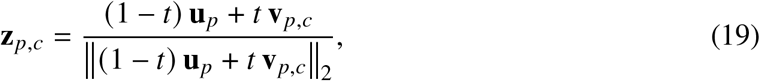

with a fixed weight *t* = 0.10. Because both projections lie in the aligned space, this step requires no further training, and it lets each protein’s representation vary across the cell types in which it appears. The representations **z**_*p,c*_ are used in all downstream analyses.

### 2.3 Training and implementation

#### SEMPER training

SEMPER couples a pretrained network encoder with a protein sequence encoder through contrastive alignment, keeping both encoders frozen and training only the projection heads. The network encoder is the multi-scale graph transformer described above, with approximately 1.4 billion trainable parameters, most of which belong to the separate GPS layers of the 442 cell types. We pretrain it once on the 442 cell type-specific PPI networks and the metagraph with the link prediction and center loss objectives (Methods 2.1.2) for 300 epochs, using GraphSAINT [165] mini-batch sampling. The encoder outputs 64-dimensional protein representations. The primary sequence encoder is ESM Cambrian 600M (`esmc_600m`, embedding dimension 1,152) [13]. For the architectural scaling analysis (Fig. 1e), we build SEMPER on six protein language model backbones in total, spanning three families, namely ESM-2 (`esm2_t6_8M_UR50D`, `esm2_t12_35M_UR50D`, `esm2_t30_150M_UR50D`) [12], ESM-3 (`esm3-sm-open-v1`, 1.4B) [175], and ESM-C (`esmc_300m`, `esmc_600m`) [13]. All back-bones share the same frozen network encoder. For each backbone, we train only the two projection heads that map the sequence and network representations into the shared 512-dimensional space, and form the final cell type-specific representations with the same interpolation (*t* = 0.10).

#### Hyperparameter tuning

We select hyperparameters for the network encoder by random search with Weights and Biases [176], choosing the configuration that maximizes the area under the receiver operating characteristic curve [177] for link prediction on the validation set. The search covers the node feature dimension (512 or 1,024), hidden dimension (8, 16, 32, or 64), output dimension (8, 16, 32, or 64), mini-batch size (8, 16, or 32), learning rate (0.0001, 0.001, or 0.01), center loss weight *λ* (0.1, 0.01, or 0.001), pooling attention channels (8, 16, or 32), number of attention heads (8, 16, or 32), dropout rate (0.5 or 0.6), and link prediction mixing coefficient *θ* (0.1 or 0.3). The selected configuration uses a node feature dimension of 512, mini-batch size of 32, hidden dimension of 16, output dimension of 8 per head with 8 attention heads, 8 pooling attention channels, learning rate of 0.001, weight decay of 0.0005, gradient clipping at a norm of 1.0, *λ* = 0.01, *θ* = 0.1, and dropout rate of 0.5.

#### Implementation details

We implement all models, data processing, and training in Python 3.8 with PyTorch 2.1.0 [178] and CUDA 11.8, and implement the network encoder with PyTorch Geometric 2.6.1 [179]. We compute sequence embeddings with the official `esm` Python package [12] and track experiments with Weights and Biases [176]. We run all computations on the Kempner Institute computing cluster with NVIDIA H100 GPUs (80 GB). Pretraining the network encoder uses Distributed Data Parallel [180] across four GPUs, whereas contrastive alignment and baseline evaluations run on a single GPU.

## 3 Benchmarking

### 3.1 Evaluation of SEMPER’s latent space

We evaluate SEMPER’s representations **z**_*p,c*_ in four analyses of the latent space.

#### 3.1.1 Evaluating cell type coherence

To test whether SEMPER’s representations retain cell type identity, we compare the similarity of proteins within and across cell types for the 30 CNS superclusters annotated by Siletti et al. [38]. For each cell type in a supercluster, we sample 100,000 protein pairs within the cell type and compute their cosine similarities, which form the intra-cell type distribution. As a null, we sample 100,000 protein pairs between that cell type and a different, randomly selected cell type. We compare the two distributions with a one-sided Mann-Whitney U test [43] and adjust for multiple testing across superclusters with the Benjamini-Hochberg procedure [44].

#### 3.1.2 Functional compactness of Gene Ontology

To test whether proteins with shared functions lie close together, we measure the compactness of Gene Ontology (GO) [45, 46] terms in SEMPER’s latent space. We represent each protein by the ℓ_2_-normalized mean of its representations across all cell types, and keep GO terms from the Biological Process, Molecular Function, and Cellular Component domains that annotate between 10 and 500 proteins. For each term, we compute a dispersion score, the mean pairwise cosine distance among the proteins it annotates [181], and compare it with the mean pairwise cosine distance across all proteins (*d*_null_ ≈ 0.93), which we use as a reference distance. We define the shift Δ = *d*_null_ − *d*_term_, so that positive values indicate terms enriched for proximity compared to average. To test significance while controlling for term sizes and overlaps between terms, we permute representations across proteins 2,000 times while keeping term memberships fixed, recompute the mean shift for each permutation, and compare the observed mean shift with this permutation distribution.

#### 3.1.3 Subcellular processes in dopaminergic neurons

To test whether SEMPER resolves functional organization within a single cell type, we analyze dopaminergic (DA) neurons (cell type 395) [47–49], identified by high expression of dopamine synthesis and vesicular transport genes (TH, SLC6A3, SLC18A2) and midbrain lineage transcription factors (EN1, LMX1A, LMX1B) [146]. We take the ℓ_2_-normalized representations of the 3,035 proteins in this cell type’s network, build a *k*-nearest-neighbor graph (*k* = 20, cosine distance), and partition it with the Leiden algorithm [182] (resolution 1.0), yielding 15 modules. We visualize the representations with UMAP (Fig. 2e). To annotate each module, we test for enrichment of GO Biological Process and Molecular Function terms against a background of the proteins in the DA neuron network and label each module with its most significantly enriched terms.

#### 3.1.4 Evaluating brain-specific protein co-abundance

To test whether SEMPER’s representations reflect brain-specific protein co-abundance, we use the tissue proteomics atlas of Laman Trip et al. [52] (Methods 1.3). For each of the 442 cell types and each of the 11 tissues, we compute the Spearman correlation between SEMPER’s cosine similarities and the tissue’s co-abundance scores over protein pairs present in both (Supplementary Figure 1). We then compare each cell type’s correlation with brain against its correlation with each other tissue using one-sided Mann-Whitney U tests with Benjamini-Hochberg correction. For the primary analysis, we focus on cell types enriched for brain-specific protein pairs, defined following Laman Trip et al. [52] as pairs whose brain co-abundance score exceeds the 95th percentile of the brain score distribution (> 0.68) and whose score in every other tissue remains below 0.5. We retain the 221 cell types in which the fraction of brain-specific pairs among atlas-covered pairs exceeds the median across all 442 cell types.

### 3.2 Protein complex membership prediction

We evaluate whether SEMPER’s representations identify cell type-specific protein complexes, framed as a zero-shot binary classification task that separates true complexes from decoys. Methods 1.4 describes the complex set, the four definitions of a cell type-specific complex, and the decoys. For a candidate complex *Q* with *m* subunits in cell type *c*, we score its cohesion as the mean pairwise cosine similarity of its subunits’ representations,]

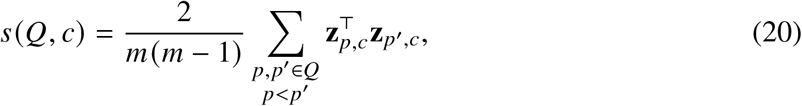

where the representations are ℓ_2_-normalized. Using *s*(*Q, c*) as a continuous score, we compute the area under the precision-recall curve (AUPRC) [183] for each cell type, separating true complexes from their matched decoys. We evaluate cell types with enough positive complexes for stable estimates, requiring at least five for the transcriptional co-expression and structural connectivity definitions and at least three for the protein co-abundance and spatial colocalization definitions, whose positive sets are sparser. This yields 349, 328, 213, and 79 cell types for the four definitions, respectively. Within each definition, we score all models on the same complexes, decoys, and cell types, and compare SEMPER with each baseline using a one-sided Wilcoxon signed-rank test [83] on paired per-cell type AUPRC values.

#### 3.2.1 Benchmarking baseline models

We compare SEMPER with three groups of models. The cell type-agnostic unimodal baselines are ESM-C [13], a sequence-only protein language model, and ORCA [57, 58], which represents each protein by graphlet degree vectors of its local topology in the cell type-agnostic PPI network. The cell type-agnostic multimodal baseline is ProTrek [59], which aligns protein sequence, three-dimensional structure (Foldseek [184]), and text-based functional annotations, and which we evaluate separately through its sequence and structure encoders (ProTrek-Seq and ProTrek-Struct). Cell type-agnostic models assign each protein a single representation, which we use in every cell type when scoring that cell type’s complexes and decoys. The cell type-specific baselines are PINNACLE [20], which learns PPI-only representations in each cell type, ORCA-CT, which computes graphlet degree vectors within each cell type-specific PPI network, and PPI-GT, SEMPER’s network encoder before sequence alignment.

## 4 Assessment of candidate drug targets for neurodegenerative disease

We evaluate whether SEMPER’s representations prioritize clinically pursued targets for five neurodegenerative diseases (Methods 1.5) in zero-shot retrieval, in which a small set of known targets serves as the query and SEMPER receives no task-specific training.

### 4.1 Zero-shot retrieval of candidate drug targets

For each disease and cell type *c*, the candidates are the proteins in the network of cell type *c*. We sample a seed set *S* of 15 known targets from those present in the network, remove the seeds from the candidates, and score every remaining candidate *p* by its mean cosine similarity to the seeds,

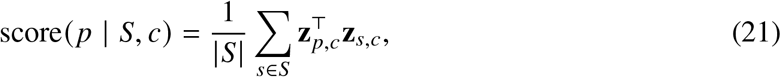

where **z**_*p,c*_ is SEMPER’s ℓ_2_-normalized representation of protein *p* in cell type *c*. We rank candidates by this score and measure Hits@*k*, the number of held-out targets among the top *k* candidates, for *k* ∈ {5, 10, 25}. We repeat the procedure for 100 independently sampled seed sets per cell type and report the mean Hits@*k* for each cell type.

#### Baselines

We compare SEMPER with three cell type-specific baselines, PINNACLE [20], PPI-GT (SEMPER’s network encoder before sequence alignment), and ORCA-CT [57, 58], and with three cell type-agnostic baselines, ProTrek-Seq, ProTrek-Struct, and ORCA. For each cell type, we restrict the cell type-agnostic baselines to the same candidates as SEMPER, and we evaluate all models on the same precomputed seed sets. We summarize each cell type-agnostic baseline by its mean Hits@*k* across cell types.

### 4.2 Comparison with a large language model

We compare SEMPER with a large language model (LLM) in the ten cell types with the highest mean SEMPER Hits@10 for each disease (Fig. 4). In each cell type, we draw 10 seed sets of 15 targets, chosen so that every target present in the network appears in at least one seed set, and evaluate all approaches on the same seed sets.

#### LLM alone

We prompt GPT-5.6-sol [90] with the seed targets and a description of the cell type, consisting of its name, class, supercluster, neurotransmitter identity, top brain regions, and marker genes, and ask for 25 additional candidate targets. Because a target can be recovered only in a cell type whose network contains it, we normalize the returned gene symbols to approved HGNC symbols and exclude proteins absent from the cell type’s network before scoring.

#### LLM scoring of SEMPER’s candidates

For each seed set, GPT-5.6-sol scores SEMPER’s top 100 candidates from 1 to 5 for their relevance as therapeutic targets for the disease in that cell type, given the same cell type description, and provides a rationale for each score. We rerank the candidates by the combined score 0.5 · score(*p* | *S, c*) + 0.5 · (*r* _*p*_ − 1)/4, where *r* _*p*_ is the LLM’s score for protein *p*.

#### Pairwise ranking with Gene Ontology annotations

We also rerank SEMPER’s top 100 candidates in a pairwise Elo tournament [185] in which the LLM compares candidates using their Gene Ontology annotations. Before the tournament, we retrieve the annotations of each candidate from the Open Targets Platform [143]. We initialize each candidate’s Elo rating between 1,100 and 1,500 in proportion to its min-max-scaled SEMPER score. In each of five rounds, we sort candidates by rating, pair adjacent candidates, and ask GPT-5.6-sol which protein of each pair is the more relevant therapeutic target for the disease in that cell type, given both proteins’ annotations and the cell type description. We update ratings after each match with a *K*-factor of 32 and rank candidates by their final ratings.

#### Gene-level comparison

To show which genes drive changes in recovery, we record the top-ranked gene for each seed set under SEMPER alone and after the tournament. For each cell type, we form the top-1 union, the set of genes ranked first in at least one of the 10 seed sets, which typically contains 4 to 10 genes. Let *A* and *B* denote the top-1 unions under SEMPER alone and after the tournament. In Fig. 6d-h, we mark genes in *B* \ *A* with a ‘+’ prefix and list *A* in a box at the SEMPER end of the slope, and we mark clinically pursued targets for the disease (Methods 1.5) with an asterisk. In each disease panel, we highlight one focal cell type, chosen as the cesssll type with the largest increase in mean Hits@25 from SEMPER to the tournament among those where SEMPER also improves on the LLM alone, or the largest increase overall if no cell type meets both conditions.

### 4.3 Architectural scaling

To test whether SEMPER’s improvement depends on a particular sequence encoder, we repeat zero-shot retrieval with six protein language model backbones from the ESM-2 [12], ESM-3 [175], and ESM-C [13] families (Methods 2). For this analysis, we use a pan-neurodegenerative target set curated from the Open Targets Platform [143] under the “neurodegenerative disease” ontology term, and compare each SEMPER variant with its sequence backbone alone.

## 5 Prediction of essential genes in dopaminergic neurons

We test whether SEMPER’s representations of dopaminergic (DA) neurons identify genes required for their survival, and whether they complement an LLM, using a genome-wide CRISPR screen in human induced pluripotent stem cell-derived midbrain DA neurons [89]. Nazeen et al. [89] performed this screen on human iPSC-derived midbrain dopaminergic neurons and identified a set of genes essential to their survival. Because these genes come from a recent experimental screen, they are less likely than literature-derived gene sets to be recoverable from text or LLM parametric knowledge alone. Because many of these essential genes are experimentally discovered and may not be historically associated with DA neurons in broad literature, this dataset serves as a rigorous benchmark to test whether a model is capturing actual physical biology rather than merely regurgitating text co-occurrences.

In each of 100 trials, we sample a seed set of 20 essential genes that are not among the 100 strongest-effect genes and compare three strategies on the same seed sets. For SEMPER alone, we rank the proteins in the DA neuron network (cell type 395) by their mean cosine similarity to the seeds, excluding the seeds, as in target retrieval (Methods 4). For the LLM alone, we prompt GPT-5.6-sol [90] with the seeds and ask for 75 additional probable essential genes, without restricting its answers to the DA neuron network. For the combination, GPT-5.6-sol scores SEMPER’s top 100 candidates from 1 to 5 for their likely essentiality and provides a rationale for each, and we rerank the candidates by 0.5 · score(*p* | *S, c*) + 0.5 · (*r* _*p*_ − 1)/4, where *r* _*p*_ is the LLM’s score for protein *p* (Methods 6). For each strategy, we measure Hits@*k*, the number of essential genes among the top *k* candidates, for *k* ∈ {25, 50, 75}, against both the 693 essential genes and the 100 strongest-effect genes.

## 6 LLM and statistical analysis

### 6.1 LLM reranking

We use GPT-5.6-sol [90] through Azure OpenAI in two ways to rerank SEMPER’s top 100 candidates, pointwise scoring and a pairwise Elo tournament.

#### Pointwise scoring

We give the LLM the seed targets, a description of the cell type, and SEMPER’s top 100 candidates with their cosine similarity scores *s*_*p*_, and ask it to return, for each candidate, a rationale and an integer relevance score *r* _*p*_ ∈ {1, … , 5} in JSON format (Supplementary Note 2). We rescale *r* _*p*_ to the unit interval and rerank candidates by

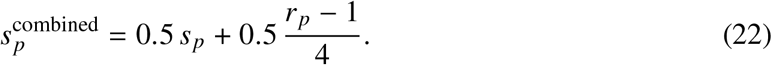

#### Pairwise Elo tournament

We rank candidates in a Swiss-system Elo tournament [185, 186] (Supplementary Note 3). Before the tournament, we map each candidate to its Ensembl gene identifier and retrieve its Gene Ontology annotations [45] from the Open Targets Platform [143], truncating each protein’s annotations to 1,500 characters. We initialize each candidate’s rating *R* between 1,100 and 1,500 in proportion to its min-max-scaled cosine similarity, so that the tournament starts from SEMPER’s ranking. In each of five rounds, we sort candidates by rating and pair adjacent candidates, so that each match compares closely ranked proteins. We send matches to the LLM in batches of three, and for each match the LLM returns, in JSON format, the winner given the two proteins’ annotations and the cell type description. The expected score of protein A against protein B is

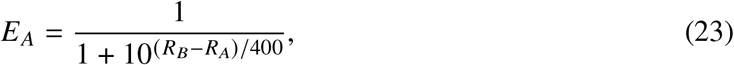

and given the outcome *S*_*A*_ ∈ {1, 0.5, 0} for a win, tie, or loss, we update A’s rating with a *K*-factor of 32 [187],

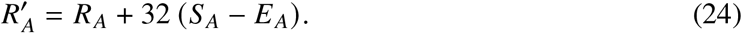

We rank candidates by their ratings after the final round.

### 6.2 Statistical analysis

#### 6.2.1 Graph clustering and manifold projection

For the analysis of dopaminergic neuron subcellular processes, local geometric relationships between cell type-specific protein embeddings were captured by constructing a *k*-nearest neighbors (*k*-NN) graph (*k* = 20) using cosine distance [188]. Unsupervised community detection was performed on this graph utilizing the Leiden algorithm [182] with a resolution parameter of 1.0 to partition the graph into discrete modules. For two-dimensional visualization of this high-dimensional manifold, Uniform Manifold Approximation and Projection (UMAP) [189] was applied to the ℓ_2_-normalized protein representations.

#### 6.2.2 Statistical testing

All statistical analyses were evaluated using a standard significance threshold of *α* = 0.05. Where applicable, raw P-values were adjusted for multiple hypothesis testing using the Benjamini-Hochberg False Discovery Rate (FDR) method [44]. Retrieval and complex analyses have a nested design, in which seed sets are sampled repeatedly within each cell type. For each cell type, we first average Hits@*k* over its 100 seed sets and then compare models with paired tests across cell types, so repeated seed sets from the same cell type are never counted as independent samples. All models are scored on the same seed sets, complexes, and decoys, so all model comparisons are paired. Where we report a mean in the text, we give the s.d. across cell types for zero-shot target retrieval and across the 100 seed trials for essential-gene prediction. We report effect sizes as differences in mean performance between models. We used nonparametric rank-based tests throughout because Hits@*k* and AUPRC are bounded and discrete; we therefore did not assume normality and did not test for it. Specific statistical tests were applied as follows:

##### Cell-type coherence

Differences between intra-cell-type and inter-cell-type cosine similarity distributions were evaluated using a one-sided Mann-Whitney U test [43], testing against the null hypothesis that intra-cell-type similarities are less than or equal to the inter-cell-type similarities.

##### GO functional compactness

The statistical significance of the GO term dispersion shift (Δ) was assessed with a label-permutation test. For each GO domain, the observed mean shift across terms was compared against a null distribution generated by randomly permuting the assignment of proteins to embeddings 2,000 times; this preserves the number of GO terms, their sizes, and their protein-overlap structure while removing any association between a protein’s embedding and its annotations. One-sided empirical *P*-values were computed as the fraction of permutations whose mean shift was at least as extreme as the observed value, and the effect size was reported as the number of standard deviations by which the observed shift exceeds the permutation null.

##### Module enrichment

Statistical overrepresentation of GO terms within identified dopaminergic neuron modules was calculated using a right-tailed hypergeometric test [190] for the GO enrichment analysis.

##### Tissue-resolved co-abundance comparison

Differences in per-cell-type Spearman correlation distributions between brain and each non-brain tissue were evaluated using one-sided Mann-Whitney U tests [43] against the null hypothesis that brain correlations are less than or equal to non-brain tissue correlations. Raw p-values were adjusted for multiple hypothesis testing across the 10 tissue comparisons using Benjamini-Hochberg FDR correction [44].

##### PPI network co-abundance validation

To validate that cell type-specific PPI networks capture biologically meaningful protein associations, the brain co-abundance scores of existing network edges were compared against those of absent protein pairs. For each cell type, absent pairs were defined as all protein pairs carrying a brain tissue co-abundance score in the atlas of Laman Trip et al. [52] whose two proteins were both present in the network but that were not network edges (up to 500,000 pairs per cell type). Brain co-abundance association scores were retrieved for every existing edge and every absent pair, and a median score was computed for each set per cell type. Across all 442 cell types, the paired per-cell-type medians (existing vs. absent) were compared using a one-sided Wilcoxon signed-rank test [83], against the null hypothesis that the existing-edge median is less than or equal to the absent-pair median.

##### Complex identification performance

To determine the statistical significance of SEMPER’s AUPRC performance gains over the baseline models across all evaluated cell types, a paired, one-sided Wilcoxon signed-rank test [83] was applied separately for each of the four definitions against each of the three cell type-specific baselines (12 comparisons; Supplementary Table 5). We report raw *P* values; Benjamini-Hochberg correction across the 12 comparisons does not change any conclusion.

##### Zero-shot target retrieval

For each disease, the per-cell-type mean Hits@10 (averaged over 100 randomly sampled seed sets) was compared between SEMPER and each cell type-specific baseline (PINNACLE, PPI-GT, ORCA-CT) using a paired, one-sided Wilcoxon signed-rank test [83], against the null hypothesis that SEMPER’s per-cell-type Hits@10 is less than or equal to the baseline’s, over the cell types expressing the disease seed set. Raw p-values were adjusted across the 15 disease-by-baseline comparisons using Benjamini-Hochberg FDR correction [44].

##### Essential-gene prediction

For each *k* ∈ {25, 50, 75}, per-trial Hits@*k* was compared across the 100 seed trials, paired by seed set, using one-sided Wilcoxon signed-rank tests [83]. We test SEMPER against the LLM, SEMPER reordered by the LLM against SEMPER, and SEMPER reordered by the LLM against the LLM, against the null hypothesis that the first method’s Hits@*k* is less than or equal to the second’s. The same tests were applied to the 693 essential genes and to the 100 genes with the strongest fitness effect. P-values were not adjusted across the three values of *k*.

##### Comparison with the LLM

Per-pair mean Hits@25 was compared between SEMPER and the LLM alone across the 50 disease and cell type pairs using a paired, one-sided Wilcoxon signed-rank test [83], against the null hypothesis that SEMPER’s Hits@25 is less than or equal to the LLM’s. Before scoring, genes nominated by the LLM that are absent from the cell type’s PPI network were removed, so that both methods rank the same candidate pool.

##### Elo tournament

Per-pair mean Hits@25 was compared between SEMPER reranked by the tournament and SEMPER alone, and between SEMPER reranked by the tournament and the LLM alone, using paired, one-sided Wilcoxon signed-rank tests [83] across all 50 disease and cell type pairs. The comparison with SEMPER alone was also tested within each disease across its 10 cell types. P-values were not adjusted across the five diseases.

