## Supplementary Information for "Multimodal protein representations across human brain cell types"

**This PDF file includes:**

Supplementary Notes 1 to 3

Supplementary Figures 1 to 3

Supplementary Tables 1 to 8

### Supplementary Note 1: LLM-based selection of CNS-relevant CORUM complexes

To build the complex-prediction benchmark for SEMPER, we use an LLM to score each human protein complex in the CORUM database [1] for relevance to the central nervous system (CNS), and retain the relevant complexes.

For each complex, the LLM (o3-mini) receives the complete CORUM record, including the complex name and synonyms, the subunit gene and protein names, the source cell line and PubMed identifiers (PMIDs), curator comments on the complex, its subunits, associated diseases and drugs, and Gene Ontology (GO) annotations. Complexes are scored in batches of five.

We prompt the model as a biomedical expert and ask it to score each complex from 1 to 5 for CNS relevance, returning JSON with a short rationale. The system and user prompts are shown below.

**System prompt**

```
You are a biomedical expert who evaluates protein COMPLEXES for
   their relevance to the central nervous system (CNS).

A high score means the complex has clear CNS/neuroscience
   relevance (neurons/glia, synapses, neurotransmission,
   neurodevelopment, neurodegeneration, myelination, blood-brain
   barrier, CNS signaling), while a low score means little or no
   CNS relevance.

Be concise, accurate, and cite cues from the provided info.
Respond ONLY with valid JSON and plain ASCII characters.
```

Prompt example (row abbreviated)

```
For each complex row below, you are given the ENTIRE ROW as
   key-value pairs (all columns), including fields like
   ComplexName, Synonyms, Organism, Cell line, PMIDs, Comments,
   GO annotations, etc.

For each row, produce an entry with:
[
  {
    "ComplexID": "STRING",      // echo the ComplexID from the row
    "Complex_name": "STRING",   // echo the complex_name from the
   row
    "rationale": "1-2 sentences explaining CNS relevance (or lack
   thereof) citing cues from the row",
    "rationale_score": 1-5
  },
  ...
]

Guidelines for rationale_score:
1 = No apparent CNS link.
2 = Weak/indirect/uncertain link.
3 = Possible CNS relevance but ambiguous.
4 = Clear CNS relevance.
5 = Strong, specific CNS role (e.g., synaptic machinery,
   neurotransmission, CNS development/degeneration).

Use only plain ASCII characters. Do not include commentary
   outside the JSON.

Here are the full rows for this batch:
[
  {
    "complex_id": "466",
    "complex_name": "Gamma-secretase complex (APH1A, PSEN1,
   PSENEN, NCSTN)",
    "synonyms": "Gamma-secretase complex",
    "organism": "Human",
    "pmid": "15286082",
    "subunits_gene_name": "APH1A;NCSTN;PSEN1;PSENEN",
    "functions_go_name": "Golgi apparatus;plasma membrane;...",
    ...
  }
]
```

Of the 5,366 human complexes, 1,425 scored 4 or 5 and were retained. We then kept the complexes whose subunits all map to proteins in the cell type-agnostic PPI network (Methods 1.2) and that have at least three distinct subunits, giving the final set of 547 complexes (Supplementary Table 4). A representative output entry for a retained complex is shown below.

Example output


```
[
  {
    "ComplexID": "row_0",
    "Complex_name": "Gamma-secretase complex (APH1A, PSEN1,
   PSENEN, NCSTN)",
    "rationale": "This complex is directly involved in the
   proteolytic processing of APP and Notch, with citations of
   amyloid beta reduction in the brain. Its role in amyloid beta
   production links it strongly to CNS pathologies like
   Alzheimer's disease.",
    "rationale_score": 5
  }
]
```

### Supplementary Note 2: LLM rescoring of SEMPER candidates

In this pipeline, a large language model scores each of SEMPER's candidate proteins for relevance to the disease context, and the score is combined with SEMPER's embedding similarity to reorder the candidates.

The large language model (GPT-5.6-sol [2]) scores candidate proteins in batches of up to $n = 100$. Given the input proteins, for example, verified essential genes for dopaminergic-neuron survival, the LLM assigns each candidate a relevance score from 1 to 5.

We combine the LLM score with SEMPER's embedding similarity into a single score per candidate. The LLM-assigned 1-5 score is rescaled to [0,1] and averaged with the cosine similarity provided by SEMPER:

$$S_{\text{combined}} = 0.5 \cdot S_{\text{similarity}} + 0.5 \cdot \left( \frac{S_{\text{rationale}} - 1}{4} \right) \tag{1}$$

where $S_{\text{similarity}}$ is the cosine similarity score computed within SEMPER's cell-type-aware latent space, and $S_{\text{rationale}}$ is the 1-5 integer score returned by the LLM. Candidates are reordered by $S_{\text{combined}}$. The LLM is given a system prompt setting its expert role, and a structured user prompt containing the batch of candidates and their similarity scores, as shown below:

**LLM System Prompt**

```
You are a biomedical expert ranking proteins for their relevance
   to a disease context.
```

Evaluator User Prompt Template

```
Given these proteins [SEED_PROTEINS] that are known essential
   genes in hPSC-derived dopamine neurons (i.e., genes for
   dopaminergic neuron survival), below is a list of proteins
   retrieved with their similarity scores to the seed proteins:

[
  {"protein": "ADCYAP1R1", "similarity": 0.550},
  {"protein": "GPR37",     "similarity": 0.543},
  {"protein": "RFX7",      "similarity": 0.556}
]

For each protein, return a JSON object like:
[
  {
    "protein": "PROTEIN_NAME",
    "rationale": "A string with 1-2 sentences that explain the
   protein's relevance, if any, to the given context",
    "rationale_score": 1-5
  },
  ...
]

Your output must use only plain ASCII characters.
```

The blocks below show a representative LLM response and the resulting combined ranking.

Raw Evaluator LLM Response


```
[
  {
    "protein": "ADCYAP1R1",
    "rationale": "ADCYAP1R1 mediates PACAP-dependent cAMP, ERK,
   and AKT signaling, which has strong neurotrophic and
   protective effects in dopaminergic neuron models.",
    "rationale_score": 5
  },
  {
    "protein": "GPR37",
    "rationale": "GPR37 is enriched in the nervous system, linked
   to parkin biology and Parkinson disease mechanisms, and can
   modulate dopaminergic neuron survival.",
    "rationale_score": 5
  },
  {
    "protein": "RFX7",
    "rationale": "RFX7 is a transcriptional regulator of stress
   and tumor-suppressive programs, but its role in dopaminergic
   neuron maintenance is poorly established.",
    "rationale_score": 2
  }
]
```


Applying the formula above to the LLM scores and SEMPER similarities gives the combined ranking:

Combined ranking


```
[
  {
    "protein": "ADCYAP1R1",
    "similarity": 0.550,
    "rationale_score": 5,
    "combined_score": 0.775
  },
  {
    "protein": "GPR37",
    "similarity": 0.543,
    "rationale_score": 5,
    "combined_score": 0.771
  },
  {
    "protein": "RFX7",
    "similarity": 0.556,
    "rationale_score": 2,
    "combined_score": 0.403
  }
]
```

### Supplementary Note 3: Pairwise LLM tournament with Gene Ontology tools

To ground the reordering in curated knowledge rather than the LLM's parametric memory alone, we reorder SEMPER's top $n = 100$ candidates with a multi-round Swiss-system Elo tournament. Each pairwise comparison is decided by a large language model from the two proteins' retrieved Gene Ontology (GO) annotations rather than from parametric knowledge alone.

Each candidate's baseline Elo is set between 1100 and 1500 by linearly scaling its SEMPER cosine similarity, so the tournament starts from SEMPER's ranking. For each candidate we retrieve GO annotations through a ToolUniverse [3] tool call, truncated to fit the context window and included in the prompt.

The tournament runs for 5 rounds. In each round, candidates are sorted by their current Elo and paired adjacently (Swiss-system), and the large language model decides each pair from the two proteins' GO annotations, selecting the more relevant target for the disease in that cell type, in batches of three. After each comparison, the winner's and loser's (or tied) ratings are updated with the standard Elo formula ($K$-factor = 32). For each comparison the model receives a single prompt combining fixed referee instructions, including the expert persona, the cell-type context, and the required output format, with that match's two proteins and their GO annotations. The instructions and an example match are shown below.

Referee instructions

```
You are an expert biomedical researcher specializing in
   Parkinson's disease.
You are refereeing a "Relevance Tournament" for disease targets
   WITHIN A SPECIFIC brain cell type:
- Cell type: Misc_165
- Cell class: NEUR
- Supercluster: Miscellaneous
- Neurotransmitter identity: VGLUT1 VGLUT2
- Predominant brain regions: Cerebral cortex: 92.1%, Hippocampus:
   5.0%, Amygdala: 2.8%
- Marker (top enriched) genes for this cell type: LINC02752,
   BARX2, IFNG-AS1, DSC3, AC116345.1, SNHG31, AC104689.2,
   AC006974.2, TMEM45B, AC072062.1

I will provide batches of 2 proteins (Protein A vs Protein B).
I have also provided their Gene Ontology (GO) annotations.
For each pair, decide: Which protein is a more functionally
   relevant and promising therapeutic target or biomarker for
   Parkinson's disease IN THIS SPECIFIC CELL TYPE -- taking its
   class, neurotransmitter identity, brain regions, and marker
   genes into account, not generic disease relevance?

### OUTPUT FORMAT (JSON ONLY):
{ "results":
  [
    {
    "match_id": 0,
    "winner": "A",
    "critique": "Brief 1-sentence reason based on the provided GO
   data."
    }
  ]
}
```

Example match input

```
### THE MATCHES AND GO DATA:
MATCH_ID 0: GRIA3 vs GABRG2
--- GRIA3 GO Data ---
{"biological_process": ["ionotropic glutamate receptor signaling
   pathway", "protein homotetramerization", "monoatomic ion
   transport"], "molecular_function": ["ligand-gated monoatomic
   ion channel activity", "protein binding"],
   "cellular_component": ["postsynaptic density membrane", "AMPA
   glutamate receptor complex", "dendritic spine", "endocytic
   vesicle membrane"]}
--- GABRG2 GO Data ---
{"biological_process": ["gamma-aminobutyric acid signaling
   pathway", "chloride transmembrane transport", "inhibitory
   synapse assembly", "monoatomic ion transport"],
   "molecular_function": ["GABA-gated chloride ion channel
   activity", "transmembrane signaling receptor activity"],
   "cellular_component": ["postsynapse", "plasma membrane"]}
```

The model returns its decision as JSON; an example comparison is shown below.

Example output

```
{
  "results": [
    {
      "match_id": 0,
      "winner": "A",
      "critique": "GRIA3 is more aligned with this VGLUT1/VGLUT2
   cortical excitatory neuron because its GO annotations directly
   implicate AMPA glutamate receptor signaling, dendritic spines,
   and postsynaptic density membranes, whereas GABRG2 primarily
   mediates inhibitory GABAergic input."
    }
  ]
}
```

Upon parsing this output, the orchestrator updates the Elo rankings. For example, if both proteins entered the match at an Elo of 1300, the victory awards GRIA3 a +16 adjustment (new Elo: 1316) while penalizing GABRG2 by −16 (new Elo: 1284). After all 5 rounds conclude, the targets are resorted by their final Elo scores, yielding the final reordered target list.

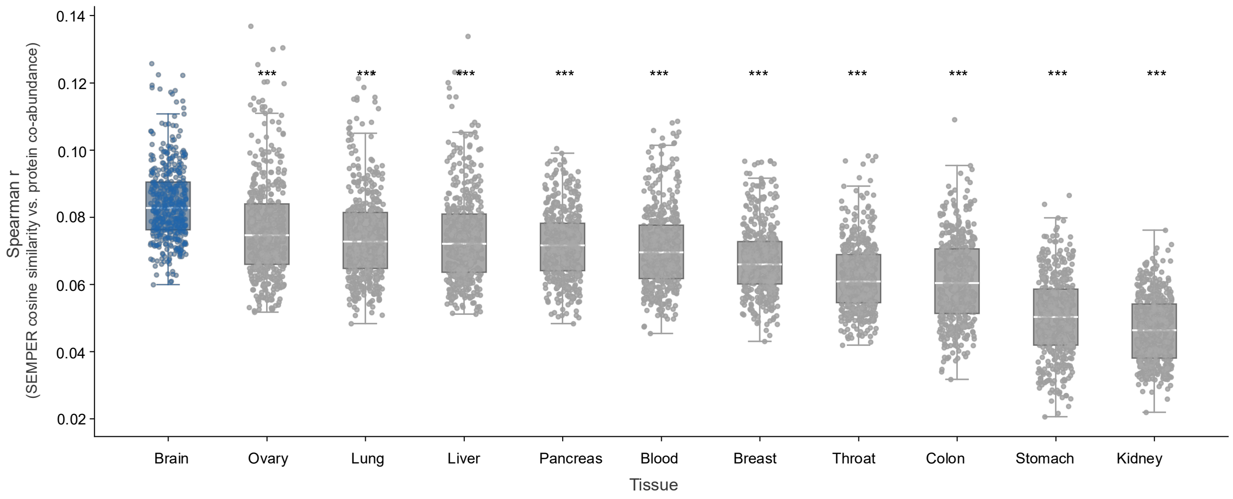


**Supplementary Figure 1: SEMPER embeddings correlate more strongly with brain protein co-abundance than with any other tissue across all cell types.** Spearman correlation between SEMPER pairwise cosine similarities and protein co-abundance scores from the Laman Trip et al. tissue proteomics atlas [4], computed per cell type across all 442 brain cell types and 11 human tissues. For each cell type, the Spearman correlation is computed across all protein pairs present in both the SEMPER embedding space and the tissue atlas. Brain co-abundance correlations (blue; median $r = 0.083$) are significantly higher than all 10 non-brain tissues (grey; BH-corrected one-sided Mann-Whitney U, $P < 0.001$ for all comparisons). Each dot represents one SEMPER cell type.

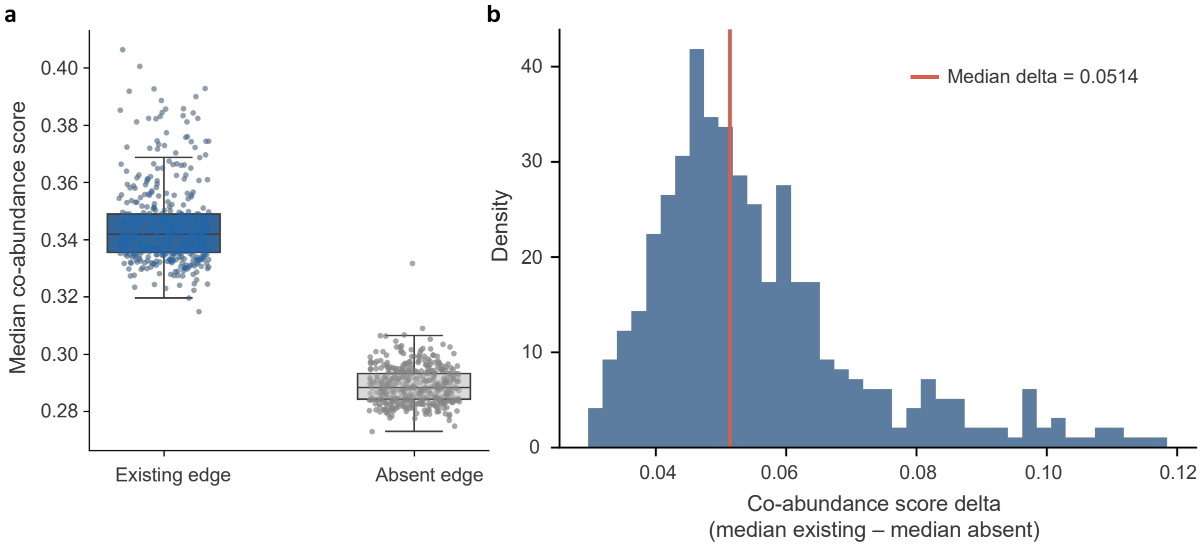


**Supplementary Figure 2: Cell type specific PPI edges capture higher brain protein co-abundance than absent protein pairs.** Validation of the cell type specific PPI networks against the brain tissue protein co-abundance atlas independently derived by Laman Trip et al. [4]. For each of the 442 cell type specific networks, existing edges were compared against absent protein pairs, defined as all atlas-scored pairs whose two proteins are both present in the network but that are not network edges, taking the first 500,000 such pairs per cell type. **(a)** Per-cell-type median brain co-abundance score for existing edges (blue) and absent pairs (grey); each dot is one cell type ($n$ = 442) and boxes denote the median and interquartile range across cell types. **(b)** Distribution across cell types of the per-cell-type co-abundance score delta (median existing − median absent); the red line marks the median delta (0.0514).

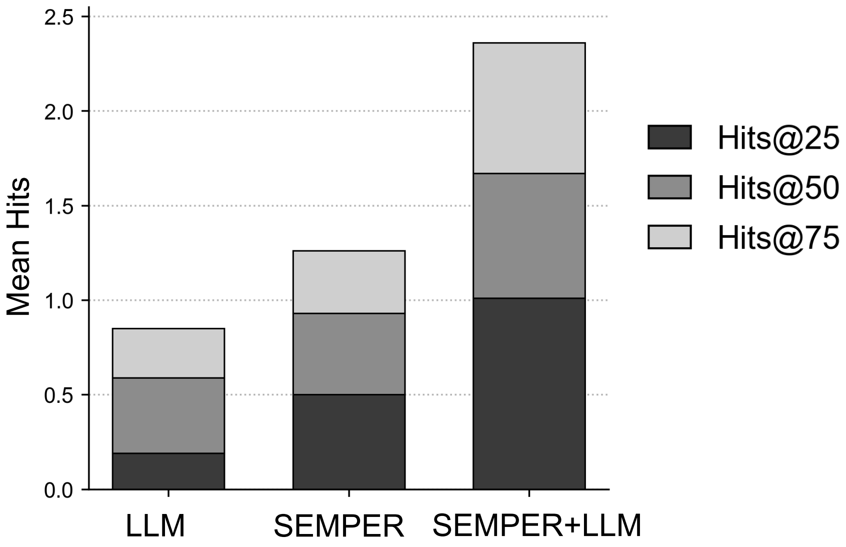


**Supplementary Figure 3: Essential-gene recovery in dopaminergic neurons on the top-100 highest-confidence subset of the screen.** Hits@K recovery of held-out essential genes restricted to the 100 genes with the strongest fitness effect in the Nazeen et al. [5] genome-wide CRISPR screen of human midbrain dopaminergic neurons. Each bar shows cumulative Hits@K (K = 25, 50, 75) for three zero-shot strategies, LLM-only (GPT-5.6-sol [2]), SEMPER, and SEMPER +LLM, averaged over 100 seed bootstraps. Mean Hits@25/50/75 are 0.19/0.59/0.85 (LLM-only), 0.50/0.93/1.26 (SEMPER), and 1.01/1.67/2.36 (SEMPER +LLM).

| **Supercluster** | ***n*** | **Mean intra** | **Mean inter** | **Shift** | $P_{\mathbf{adj}}$ |
|---|---|---|---|---|---|
| Miscellaneous | 22 | 0.0895 | 0.0581 | 0.0314 | $< 10^{-300}$ |
| Vascular | 11 | 0.0805 | 0.0502 | 0.0303 | $< 10^{-300}$ |
| Microglia | 9 | 0.0753 | 0.0482 | 0.0270 | $< 10^{-300}$ |
| Upper-layer intratelencephalic | 16 | 0.0849 | 0.0603 | 0.0246 | $< 10^{-300}$ |
| Fibroblast | 6 | 0.0816 | 0.0598 | 0.0217 | $< 10^{-300}$ |
| Astrocyte | 13 | 0.0800 | 0.0588 | 0.0212 | $< 10^{-300}$ |
| MGE interneuron | 26 | 0.0809 | 0.0600 | 0.0210 | $< 10^{-300}$ |
| Ependymal | 7 | 0.0774 | 0.0572 | 0.0203 | $< 10^{-300}$ |
| Hippocampal dentate gyrus | 7 | 0.0731 | 0.0534 | 0.0197 | $< 10^{-300}$ |
| Deep-layer intratelencephalic | 14 | 0.0818 | 0.0629 | 0.0190 | $< 10^{-300}$ |
| Committed oligodendrocyte precursor | 7 | 0.0699 | 0.0509 | 0.0189 | $< 10^{-300}$ |
| Medium spiny neuron | 18 | 0.0807 | 0.0627 | 0.0180 | $< 10^{-300}$ |
| Amygdala excitatory | 18 | 0.0807 | 0.0633 | 0.0174 | $< 10^{-300}$ |
| Thalamic excitatory | 17 | 0.0702 | 0.0537 | 0.0165 | $< 10^{-300}$ |
| Cerebellar inhibitory | 10 | 0.0727 | 0.0562 | 0.0164 | $< 10^{-300}$ |
| Oligodendrocyte | 8 | 0.0723 | 0.0561 | 0.0162 | $< 10^{-300}$ |
| Choroid plexus | 5 | 0.0791 | 0.0629 | 0.0162 | $< 10^{-300}$ |
| Eccentric medium spiny neuron | 14 | 0.0730 | 0.0572 | 0.0158 | $< 10^{-300}$ |
| Deep-layer corticothalamic and 6b | 17 | 0.0768 | 0.0615 | 0.0153 | $< 10^{-300}$ |
| CGE interneuron | 21 | 0.0678 | 0.0527 | 0.0150 | $< 10^{-300}$ |
| LAMP5-LHX6 and Chandelier | 12 | 0.0759 | 0.0609 | 0.0149 | $< 10^{-300}$ |
| Oligodendrocyte precursor | 5 | 0.0730 | 0.0600 | 0.0129 | $6.47 \times 10^{-244}$ |
| Deep-layer near-projecting | 13 | 0.0735 | 0.0606 | 0.0129 | $< 10^{-300}$ |
| Hippocampal CA1-3 | 14 | 0.0771 | 0.0644 | 0.0127 | $< 10^{-300}$ |
| Mammillary body | 9 | 0.0694 | 0.0569 | 0.0126 | $< 10^{-300}$ |
| Splatter | 90 | 0.0594 | 0.0473 | 0.0121 | $< 10^{-300}$ |
| Hippocampal CA4 | 9 | 0.0631 | 0.0511 | 0.0121 | $< 10^{-300}$ |
| Upper rhombic lip | 6 | 0.0624 | 0.0511 | 0.0113 | $2.53 \times 10^{-219}$ |
| Lower rhombic lip | 6 | 0.0623 | 0.0535 | 0.0088 | $1.08 \times 10^{-124}$ |
| Midbrain-derived inhibitory | 12 | 0.0660 | 0.0575 | 0.0085 | $8.19 \times 10^{-252}$ |

**Supplementary Table 1: Per-supercluster intra- vs. inter-cell-type embedding similarity.** For each of 30 CNS superclusters, mean pairwise cosine similarity between SEMPER protein embeddings was computed within the same cell type (Mean intra) and between distinct cell types (Mean inter), each over $10^5$ randomly sampled protein pairs per cell type (Methods). $n$ indicates the number of cell types in each supercluster. Shift is the difference between Mean intra and Mean inter. Statistical significance was assessed using a one-sided Mann-Whitney U test [6]; reported $P_{\text{adj}}$ values are Benjamini-Hochberg FDR-corrected [7] across all superclusters. All comparisons reach $P_{\text{adj}} < 10^{-100}$. Rows are ordered by descending shift magnitude.

**Supplementary Table 2: Top GO term enrichments per dopaminergic embedding cluster.** For each Leiden cluster identified in the dopaminergic (DA) neuron embedding subspace (Fig. 2e), up to five most-significant Gene Ontology terms per domain (Biological Process, BP; Molecular Function, MF) are reported. Enrichments were computed against the DA-expressed proteome background ($N$ = 3,035) using a one-sided hypergeometric test, with Benjamini-Hochberg FDR correction [7]. Only terms reaching $P_{\text{adj}} < 0.05$ are shown. Cluster annotations summarize the dominant pathway theme; cluster 6 yielded no FDR-significant terms in either domain. Columns: $k$ = cluster genes annotated to the GO term; $K$ = total background genes annotated to the term; Fold = enrichment score (observed / expected ratio).

| Cluster | Annotation | Domain | GO ID | GO term | Fold | $k$ / $K$ | $P_{\text{adj}}$ |
|---|---|---|---|---|---|---|---|
| 0 | Neuropeptide / TF activity | MF | GO:0004222 | metalloendopeptidase activity | 5.27 | 6/8 | 0.026 |
| | | MF | GO:0001227 | DNA-binding transcription repressor activity, RNA polymerase II-specific | 2.69 | 13/34 | 0.033 |
| 1 | Cyclic nucleotide signaling | MF | GO:0004115 | 3',5'-cyclic-AMP phosphodiesterase activity | 5.58 | 6/8 | 0.018 |
| | | MF | GO:0008429 | phosphatidylethanolamine binding | 6.20 | 5/6 | 0.018 |
| | | MF | GO:0047555 | 3',5'-cyclic-GMP phosphodiesterase activity | 5.31 | 5/7 | 0.038 |
| 2 | Ribosome & translation | BP | GO:0032543 | mitochondrial translation | 5.60 | 19/28 | $2.25 \times 10^{-9}$ |
| | | BP | GO:0006412 | translation | 3.19 | 17/44 | 0.001 |
| | | BP | GO:0006614 | SRP-dependent cotranslational protein targeting to membrane | 6.87 | 5/6 | 0.017 |
| | | MF | GO:0003735 | structural constituent of ribosome | 4.95 | 24/40 | $7.26 \times 10^{-11}$ |
| | | MF | GO:0003723 | RNA binding | 2.45 | 55/185 | $1.81 \times 10^{-9}$ |
| | | MF | GO:0003743 | translation initiation factor activity | 4.81 | 7/12 | 0.006 |
| | | MF | GO:0003756 | protein disulfide isomerase activity | 6.87 | 5/6 | 0.006 |
| | | MF | GO:0008312 | 7S RNA binding | 8.25 | 4/4 | 0.006 |
| 3 | Excitatory synapse | BP | GO:0035249 | synaptic transmission, glutamatergic | 6.43 | 9/15 | $3.29 \times 10^{-4}$ |
| | | BP | GO:0035235 | ionotropic glutamate receptor signaling pathway | 8.34 | 7/9 | $3.29 \times 10^{-4}$ |
| | | BP | GO:0016339 | calcium-dependent cell-cell adhesion | 9.19 | 6/7 | $5.08 \times 10^{-4}$ |
| | | BP | GO:0007043 | cell-cell junction assembly | 7.66 | 5/7 | 0.012 |
| | | BP | GO:0044331 | cell-cell adhesion mediated by cadherin | 5.85 | 6/11 | 0.015 |
| | | MF | GO:0005245 | voltage-gated calcium channel activity | 8.04 | 6/8 | 0.002 |
| | | MF | GO:1904315 | transmitter-gated monoatomic ion channel activity involved in regulation of postsynaptic membrane potential | 6.26 | 7/12 | 0.002 |
| | | MF | GO:0004971 | AMPA glutamate receptor activity | 10.72 | 4/4 | 0.003 |
| | | MF | GO:0022849 | glutamate-gated calcium ion channel activity | 8.58 | 4/5 | 0.012 |
| | | MF | GO:0003777 | microtubule motor activity | 5.96 | 5/9 | 0.017 |

*Continued on next page.*

*Table 2 continued from previous page.*

| Cluster | Annotation | Domain | GO ID | GO term | Fold | $k$ / $K$ | $P_{adj}$ |
|---|---|---|---|---|---|---|---|
| 4 | Ubiquitin-proteasome | BP | GO:0043161 | proteasome-mediated ubiquitin-dependent protein catabolic process | 3.58 | 22/74 | $1.74 \times 10^{-5}$ |
| | | BP | GO:0060337 | type I interferon-mediated signaling pathway | 10.04 | 5/6 | 0.004 |
| | | BP | GO:0033209 | tumor necrosis factor-mediated signaling pathway | 6.57 | 6/11 | 0.012 |
| | | BP | GO:0045116 | protein neddylation | 9.63 | 4/5 | 0.019 |
| | | MF | GO:0061630 | ubiquitin protein ligase activity | 3.01 | 16/64 | 0.005 |
| | | MF | GO:0008270 | zinc ion binding | 1.68 | 41/294 | 0.024 |
| | | MF | GO:0004298 | threonine-type endopeptidase activity | 12.04 | 3/3 | 0.024 |
| | | MF | GO:0003720 | telomerase activity | 12.04 | 2/2 | 0.048 |
| | | MF | GO:0030229 | very-low-density lipoprotein particle receptor activity | 12.04 | 2/2 | 0.048 |
| 5 | Inhibitory synapse | BP | GO:0007214 | gamma-aminobutyric acid signaling pathway | 9.63 | 6/9 | $7.07 \times 10^{-4}$ |
| | | BP | GO:1902476 | chloride transmembrane transport | 7.23 | 8/16 | $7.07 \times 10^{-4}$ |
| | | BP | GO:0051932 | synaptic transmission, GABAergic | 8.67 | 6/10 | 0.001 |
| | | BP | GO:1904862 | inhibitory synapse assembly | 9.03 | 5/8 | 0.003 |
| | | BP | GO:0010880 | regulation of release of sequestered calcium ion into cytosol by sarcoplasmic reticulum | 14.45 | 3/3 | 0.011 |
| | | MF | GO:0022851 | GABA-gated chloride ion channel activity | 12.39 | 6/7 | $2.40 \times 10^{-5}$ |
| | | MF | GO:0004890 | GABA-A receptor activity | 12.39 | 6/7 | $2.40 \times 10^{-5}$ |
| | | MF | GO:0008503 | benzodiazepine receptor activity | 14.45 | 5/5 | $3.59 \times 10^{-5}$ |
| 7 | Mitotic / cytoskeleton | BP | GO:0000278 | mitotic cell cycle | 5.78 | 9/27 | 0.002 |
| | | MF | GO:1990890 | netrin receptor binding | 17.34 | 2/2 | 0.026 |
| | | MF | GO:0001642 | group III metabotropic glutamate receptor activity | 17.34 | 2/2 | 0.026 |
| | | MF | GO:0047389 | glycerophosphocholine phosphodiesterase activity | 17.34 | 2/2 | 0.026 |
| | | MF | GO:0004438 | phosphatidylinositol-3-phosphate phosphatase activity | 8.67 | 3/6 | 0.026 |
| | | MF | GO:0019901 | protein kinase binding | 2.54 | 12/82 | 0.026 |
| 8 | GPCR / ERK signaling | BP | GO:0070374 | positive regulation of ERK1 and ERK2 cascade | 5.64 | 11/35 | $5.71 \times 10^{-4}$ |
| | | BP | GO:0042060 | wound healing | 8.98 | 6/12 | 0.003 |
| | | BP | GO:0007189 | adenylate cyclase-activating G protein-coupled receptor signaling pathway | 5.99 | 8/24 | 0.003 |
| | | BP | GO:0001525 | angiogenesis | 3.80 | 11/52 | 0.008 |
| | | BP | GO:0050919 | negative chemotaxis | 8.98 | 5/10 | 0.008 |
| | | MF | GO:0031683 | G-protein beta/gamma-subunit complex binding | 8.98 | 5/10 | 0.011 |

*Continued on next page.*

*Table 2 continued from previous page.*

| Cluster | Annotation | Domain | GO ID | GO term | Fold | $k$ / $K$ | $P_{adj}$ |
|---|---|---|---|---|---|---|---|
| | | MF | GO:0005104 | fibroblast growth factor receptor binding | 13.47 | 3/4 | 0.023 |
| | | MF | GO:0035255 | ionotropic glutamate receptor binding | 8.98 | 4/8 | 0.023 |
| | | MF | GO:0031752 | D5 dopamine receptor binding | 17.96 | 2/2 | 0.036 |
| | | MF | GO:0090722 | receptor-receptor interaction | 17.96 | 2/2 | 0.036 |
| 9 | Chromatin remodeling | MF | GO:0042826 | histone deacetylase binding | 5.11 | 6/23 | 0.017 |
| | | MF | GO:0003713 | transcription coactivator activity | 3.52 | 9/50 | 0.017 |
| | | MF | GO:0140297 | DNA-binding transcription factor binding | 5.59 | 6/21 | 0.017 |
| | | MF | GO:1990782 | protein tyrosine kinase binding | 9.79 | 3/6 | 0.038 |
| 10 | Autophagy & lysosome | BP | GO:0016241 | regulation of macroautophagy | 10.05 | 12/25 | $6.07 \times 10^{-8}$ |
| | | BP | GO:0048388 | endosomal lumen acidification | 18.31 | 7/8 | $4.19 \times 10^{-7}$ |
| | | BP | GO:0007042 | lysosomal lumen acidification | 14.65 | 7/10 | $3.86 \times 10^{-6}$ |
| | | BP | GO:0061795 | Golgi lumen acidification | 17.94 | 6/7 | $4.00 \times 10^{-6}$ |
| | | BP | GO:0097401 | synaptic vesicle lumen acidification | 15.70 | 6/8 | $1.02 \times 10^{-5}$ |
| | | MF | GO:0046961 | proton-transporting ATPase activity, rotational mechanism | 16.10 | 10/13 | $9.48 \times 10^{-10}$ |
| | | MF | GO:0046933 | proton-transporting ATP synthase activity, rotational mechanism | 8.97 | 6/14 | $9.62 \times 10^{-4}$ |
| | | MF | GO:0004862 | cAMP-dependent protein kinase inhibitor activity | 13.95 | 4/6 | 0.002 |
| | | MF | GO:0034236 | protein kinase A catalytic subunit binding | 10.47 | 4/8 | 0.006 |
| | | MF | GO:0008379 | thioredoxin peroxidase activity | 15.70 | 3/4 | 0.007 |
| 11 | Mitochondrial respiration | BP | GO:1902600 | proton transmembrane transport | 9.56 | 30/75 | $1.70 \times 10^{-21}$ |
| | | BP | GO:0042776 | proton motive force-driven mitochondrial ATP synthesis | 13.04 | 24/44 | $2.98 \times 10^{-21}$ |
| | | BP | GO:0006120 | mitochondrial electron transport, NADH to ubiquinone | 15.93 | 20/30 | $3.12 \times 10^{-20}$ |
| | | BP | GO:0006119 | oxidative phosphorylation | 19.55 | 9/11 | $2.82 \times 10^{-10}$ |
| | | BP | GO:0006123 | mitochondrial electron transport, cytochrome c to oxygen | 15.93 | 8/12 | $4.01 \times 10^{-8}$ |
| | | MF | GO:0008137 | NADH dehydrogenase (ubiquinone) activity | 17.07 | 20/28 | $6.04 \times 10^{-21}$ |
| | | MF | GO:0004129 | cytochrome-c oxidase activity | 17.92 | 6/8 | $2.69 \times 10^{-6}$ |
| | | MF | GO:0009055 | electron transfer activity | 10.24 | 6/14 | $1.56 \times 10^{-4}$ |
| | | MF | GO:0001758 | retinal dehydrogenase (NAD+) activity | 23.90 | 3/3 | $7.70 \times 10^{-4}$ |
| | | MF | GO:0050660 | flavin adenine dinucleotide binding | 10.62 | 4/9 | 0.002 |
| 12 | Viral / amyloid metabolism | BP | GO:0045070 | positive regulation of viral genome replication | 14.78 | 3/8 | 0.020 |

*Continued on next page.*

*Table 2 continued from previous page.*

| Cluster | Annotation | Domain | GO ID | GO term | Fold | $k$ / $K$ | $P_{adj}$ |
|---|---|---|---|---|---|---|---|
| | | BP | GO:0044321 | response to leptin | 39.42 | 2/2 | 0.020 |
| | | BP | GO:0042982 | amyloid precursor protein metabolic process | 39.42 | 2/2 | 0.020 |
| | | BP | GO:0060216 | definitive hemopoiesis | 26.28 | 2/3 | 0.035 |
| | | BP | GO:0009791 | post-embryonic development | 9.10 | 3/13 | 0.046 |
| 13 | Transcription regulation | BP | GO:0006357 | regulation of transcription by RNA polymerase II | 12.39 | 10/79 | $2.97 \times 10^{-8}$ |
| | | BP | GO:0070979 | protein K11-linked ubiquitination | 32.63 | 2/6 | 0.010 |
| | | BP | GO:0007595 | lactation | 19.58 | 2/10 | 0.019 |
| | | BP | GO:0000122 | negative regulation of transcription by RNA polymerase II | 3.79 | 5/129 | 0.029 |
| | | BP | GO:0007005 | mitochondrion organization | 9.79 | 2/20 | 0.035 |
| | | MF | GO:0008270 | zinc ion binding | 5.99 | 18/294 | $2.66 \times 10^{-10}$ |
| | | MF | GO:0000981 | DNA-binding transcription factor activity, RNA polymerase II-specific | 18.36 | 9/48 | $2.64 \times 10^{-9}$ |
| | | MF | GO:0000978 | RNA polymerase II cis-regulatory region sequence-specific DNA binding | 8.06 | 7/85 | $6.43 \times 10^{-5}$ |
| | | MF | GO:0003677 | DNA binding | 5.90 | 5/83 | 0.004 |
| | | MF | GO:0061631 | ubiquitin conjugating enzyme activity | 21.76 | 2/9 | 0.008 |
| 14 | Axon guidance | BP | GO:2001222 | regulation of neuron migration | 51.01 | 2/7 | 0.003 |
| | | BP | GO:0038007 | netrin-activated signaling pathway | 59.51 | 2/6 | 0.003 |
| | | BP | GO:0007409 | axonogenesis | 13.73 | 3/39 | 0.004 |
| | | BP | GO:0060291 | long-term synaptic potentiation | 32.46 | 2/11 | 0.004 |
| | | BP | GO:0007613 | memory | 22.32 | 2/16 | 0.007 |
| | | MF | GO:0008201 | heparin binding | 12.75 | 2/28 | 0.041 |

| Tissue | Median $r$ | Mean $r$ | Δ median | MW $U$ | $p_{\text{raw}}$ | $p_{\text{BH}}$ |
|---|---|---|---|---|---|---|
| **All clusters** (brain reference: median $r$ = 0.0817, mean $r$ = 0.0831, $n$ = 442 cell types per group) | | | | | | |
| Ovary | 0.0743 | 0.0762 | 0.0074 | 131,697 | $1.6 \times 10^{-19}$ | $1.6 \times 10^{-19}$ |
| Lung | 0.0723 | 0.0741 | 0.0094 | 140,550 | $7.1 \times 10^{-30}$ | $7.9 \times 10^{-30}$ |
| Pancreas | 0.0715 | 0.0715 | 0.0101 | 151,058 | $3.3 \times 10^{-45}$ | $4.7 \times 10^{-45}$ |
| Liver | 0.0705 | 0.0722 | 0.0112 | 147,268 | $2.7 \times 10^{-39}$ | $3.3 \times 10^{-39}$ |
| Blood | 0.0686 | 0.0692 | 0.0131 | 157,722 | $1.2 \times 10^{-56}$ | $2.0 \times 10^{-56}$ |
| Breast | 0.0657 | 0.0668 | 0.0160 | 168,569 | $4.0 \times 10^{-78}$ | $7.9 \times 10^{-78}$ |
| Throat | 0.0604 | 0.0614 | 0.0212 | 180,744 | $1.9 \times 10^{-106}$ | $6.4 \times 10^{-106}$ |
| Colon | 0.0596 | 0.0600 | 0.0221 | 178,805 | $1.2 \times 10^{-101}$ | $3.1 \times 10^{-101}$ |
| Stomach | 0.0464 | 0.0465 | 0.0353 | 193,696 | $1.8 \times 10^{-141}$ | $9.1 \times 10^{-141}$ |
| Kidney | 0.0443 | 0.0441 | 0.0374 | 195,154 | $1.0 \times 10^{-145}$ | $1.0 \times 10^{-144}$ |
| **Brain-enriched clusters** (brain reference: median $r$ = 0.0832, mean $r$ = 0.0837, $n$ = 221 cell types per group) | | | | | | |
| Ovary | 0.0687 | 0.0704 | 0.0144 | 39,376 | $4.1 \times 10^{-29}$ | $4.1 \times 10^{-29}$ |
| Lung | 0.0657 | 0.0675 | 0.0174 | 41,982 | $2.2 \times 10^{-39}$ | $2.4 \times 10^{-39}$ |
| Pancreas | 0.0655 | 0.0664 | 0.0177 | 43,392 | $1.3 \times 10^{-45}$ | $1.8 \times 10^{-45}$ |
| Liver | 0.0638 | 0.0665 | 0.0193 | 42,385 | $4.1 \times 10^{-41}$ | $5.1 \times 10^{-41}$ |
| Blood | 0.0634 | 0.0641 | 0.0198 | 44,305 | $6.5 \times 10^{-50}$ | $1.1 \times 10^{-49}$ |
| Breast | 0.0617 | 0.0619 | 0.0215 | 46,171 | $2.6 \times 10^{-59}$ | $5.2 \times 10^{-59}$ |
| Throat | 0.0554 | 0.0569 | 0.0278 | 47,286 | $2.5 \times 10^{-65}$ | $8.5 \times 10^{-65}$ |
| Colon | 0.0531 | 0.0542 | 0.0300 | 47,218 | $6.0 \times 10^{-65}$ | $1.5 \times 10^{-64}$ |
| Stomach | 0.0412 | 0.0423 | 0.0419 | 48,658 | $4.0 \times 10^{-73}$ | $2.0 \times 10^{-72}$ |
| Kidney | 0.0379 | 0.0393 | 0.0453 | 48,835 | $3.6 \times 10^{-74}$ | $3.6 \times 10^{-73}$ |

**Supplementary Table 3: Per-cell-type Spearman correlations between SEMPER embedding similarity and tissue protein co-abundance, by tissue.** Spearman correlation between SEMPER embedding pairwise cosine similarities and protein co-abundance scores from the Laman Trip et al. tissue proteomics atlas [4], computed per cell type across 11 human tissues. For each tissue, the per-cell-type correlations ($r$) were compared against brain using a one-sided Mann-Whitney $U$ test (alternative: brain > tissue). Median $r$, mean $r$, and the brain-minus-tissue difference in median $r$ (Δ median) are reported per tissue. The upper section includes all 442 SEMPER cell types; the lower section is restricted to cell types where the fraction of brain-specific protein pairs, defined as pairs with brain co-abundance score > 0.68 (95th percentile) and maximum score across all other tissues < 0.5, exceeds the median across all 442 cell types ($n$ = 221). MW $U$ is the Mann-Whitney rank-sum statistic and $p_{\text{BH}}$ is the BH-adjusted $p$-value across the ten tissues within each subset.

**Supplementary Table 4: CNS-relevant CORUM protein complexes.** Provided as a separate file (`SI_Table4_complexes.xlsx`). Lists the $n$ = 547 CNS-relevant human protein complexes curated from CORUM [1]. Each complex has an LLM relevance score of at least 4 and at least three distinct subunits, all present in the cell type-agnostic PPI network. Subunit counts range from 3 to 27. The columns are described below.

**`complex_id`** CORUM complex identifier.

**`complex_name`** CORUM complex name.

**`pmid`** PubMed ID of the supporting reference.

**`subunits_gene_name`** Semicolon-delimited member gene symbols.

**`subunits_protein_name`** Semicolon-delimited member protein names.

**`functions_go_id`** Semicolon-delimited Gene Ontology term IDs.

**`functions_go_name`** Semicolon-delimited Gene Ontology term names.

**`functions_go_ontology`** GO domain for each term (molecular_function, cellular_component, or biological_process).

**`subunits_ensp`** Semicolon-delimited Ensembl protein (ENSP) identifiers.

| **Protein Complex Definition** | **Baseline** | ***N*** | ***P*** |
| --- | --- | --- | --- |
| (1) Transcriptional co-expression | PINNACLE | 349 | $3.30 \times 10^{-59}$ |
| | PPI-GT | 349 | $7.87 \times 10^{-59}$ |
| | ORCA-CT | 349 | $3.01 \times 10^{-59}$ |
| (2) Structural connectivity | PINNACLE | 328 | $1.05 \times 10^{-55}$ |
| | PPI-GT | 328 | $2.17 \times 10^{-55}$ |
| | ORCA-CT | 328 | $8.24 \times 10^{-56}$ |
| (3) Protein co-abundance | PINNACLE | 213 | $7.62 \times 10^{-37}$ |
| | PPI-GT | 213 | $2.21 \times 10^{-36}$ |
| | ORCA-CT | 213 | $7.25 \times 10^{-37}$ |
| (4) Spatial colocalization | PINNACLE | 79 | $6.45 \times 10^{-15}$ |
| | PPI-GT | 79 | $1.71 \times 10^{-13}$ |
| | ORCA-CT | 79 | $7.51 \times 10^{-15}$ |

**Supplementary Table 5: Statistical evaluation of cell type specific protein complex recovery performance.** Paired, one-sided Wilcoxon signed-rank tests comparing per-cell-type AUPRC distributions for SEMPER against three context-aware baselines (PINNACLE, PPI-GT, and ORCA-CT) across four distinct macromolecular complex definitions. The number of paired cell types ($N$) is the number of cell types with at least five positive complexes (transcriptional co-expression and structural connectivity) or at least three positive complexes (protein co-abundance and spatial colocalization). Reported $p$-values evaluate the specific alternative hypothesis that SEMPER yields higher AUPRC metrics than the respective baseline, corresponding directly to the distributions plotted in Fig. 3b-e. Benjamini-Hochberg FDR correction across all 12 comparisons does not alter any statistical conclusions.

**Supplementary Table 6: Clinically pursued therapeutic targets for five neurodegenerative diseases.** Provided as a separate multi-sheet workbook (`SI_Table6_targets.xlsx`). Ground-truth targets were retrieved from the Open Targets Platform [8] and strictly filtered to protein-coding genes targeted by drugs or compounds in completed Phase 2 trials or in Phase 3 or 4 trials of any status, then mapped from HGNC nomenclature [9] to SEMPER's protein entities. The first sheet (`Table of Contents`) describes every sheet. The five subsequent sheets give the per-disease target pools, one row per target gene: Parkinson's disease (`PD - Target Genes Summarized`, 97 genes), Huntington's disease (`HD - Target Genes Summarized`, 86 genes), multiple sclerosis (`MS - Target Genes Summarized`, 166 genes), amyotrophic lateral sclerosis (`ALS - Target Genes Summarized`, 98 genes), and Alzheimer's disease (`AD - Target Genes Summarized`, 130 genes). All five disease sheets share an identical column layout:

**`diseaseId`** MONDO/EFO disease ontology identifier.

**`targetId_genename`** HGNC gene symbol of the target.

**`targetId`** Ensembl gene (ENSG) identifier.

**`targetFromSourceId`** UniProt protein accession, or the Ensembl gene identifier for the two antisense oligonucleotide targets (*HTT* and *SOD1*).

**`Max Completed Clinical Phase`** Highest clinical phase reached by any associated drug or compound. Phase 2 counts only if completed, and Phase 3 and 4 count at any trial status.

**`Associated drugId(s)`** Aggregated ChEMBL identifiers of associated drugs/compounds.

**`drugbank`** Corresponding DrugBank identifiers, where available.

**`Target/Enzyme/Transporter`** DrugBank functional role of the protein, where annotated.

**`Pharmacological action`** Mechanism of the drug-target interaction, where annotated.

**`Therapeutic Intent`** Manually curated label of whether the associated agents are disease-modifying, symptomatic, or both.

**`Notes`** Free-text curation notes.

| Disease | Baseline | $N$ | $P$ |
|---|---|---|---|
| Parkinson's disease | PINNACLE | 346 | $1.29 \times 10^{-58}$ |
| | PPI-GT | 346 | $1.09 \times 10^{-58}$ |
| | ORCA-CT | 346 | $9.13 \times 10^{-59}$ |
| Alzheimer's disease | PINNACLE | 401 | $5.46 \times 10^{-64}$ |
| | PPI-GT | 401 | $7.49 \times 10^{-67}$ |
| | ORCA-CT | 401 | $2.10 \times 10^{-65}$ |
| Multiple sclerosis | PINNACLE | 404 | $8.01 \times 10^{-64}$ |
| | PPI-GT | 404 | $6.55 \times 10^{-66}$ |
| | ORCA-CT | 404 | $1.38 \times 10^{-58}$ |
| Amyotrophic lateral sclerosis | PINNACLE | 395 | $6.13 \times 10^{-66}$ |
| | PPI-GT | 395 | $3.13 \times 10^{-66}$ |
| | ORCA-CT | 395 | $2.20 \times 10^{-64}$ |
| Huntington's disease | PINNACLE | 129 | $9.40 \times 10^{-23}$ |
| | PPI-GT | 129 | $2.10 \times 10^{-22}$ |
| | ORCA-CT | 129 | $1.01 \times 10^{-22}$ |

**Supplementary Table 7: Statistical evaluation of zero-shot target retrieval performance.** Paired, one-sided Wilcoxon signed-rank tests [10] comparing per-cell-type Hits@10 distributions for SEMPER against three context-aware baselines (PINNACLE [11], the PPI Graph Transformer, and cell type specific graphlet signatures via ORCA-CT [12, 13]) across five neurodegenerative diseases. The number of paired cell types ($N$) reflects the subset of cell type specific networks expressing the required disease-relevant seed proteins. All comparisons reach $P < 10^{-21}$, indicating that SEMPER's per-cell-type retrieval consistently exceeds context-aware baselines across all diseases. Reported $p$-values are raw; Benjamini-Hochberg FDR correction across the 15 comparisons does not alter any conclusions.

| Gene | LLM-only | SEMPER | SEMPER+LLM |
|---|---|---|---|
| MZT2A | ✓ | ✓ | ✓ |
| GCHFR | | ✓ | ✓ |
| PRKAA1 | | ✓ | ✓ |
| HSP90AA1 | ✓ | | |
| MTHFD1L | ✓ | | |
| PSMB5 | ✓ | | |
| SNAP25 | ✓ | | |
| TIMM44 | ✓ | | |
| TIMM8B | ✓ | | |
| ARHGEF3 | | ✓ | |
| EEF1B2 | | ✓ | |
| FBXO2 | | ✓ | |
| GNAS | | ✓ | |
| GNG3 | | ✓ | |
| HMG20B | | ✓ | |
| MZT2B | | ✓ | |
| NDFIP2 | | ✓ | |
| RALGPS1 | | ✓ | |
| RAPGEF6 | | ✓ | |
| UQCR10 | | ✓ | |
| CHORDC1 | | | ✓ |
| DTNBP1 | | | ✓ |
| HTT | | | ✓ |
| NAPB | | | ✓ |
| STMN2 | | | ✓ |
| TTC19 | | | ✓ |

**Supplementary Table 8: DA-essential genes recovered by each retrieval paradigm.** Each row is a gene from the genome-wide Nazeen et al. [5] DA-neuron essentiality screen that was pooled into the top-3 candidates of at least one of three paradigms across the 100 seed sets: LLM-only (parametric reasoning by GPT-5.6-sol [2]); SEMPER (zero-shot embedding retrieval on the cell type resolved dopaminergic-neuron representation); SEMPER +LLM (LLM-evaluator reranking of SEMPER's top-100 candidates). Checkmarks indicate which paradigms recovered the gene; pooled recovery is 7 (LLM-only), 14 (SEMPER), and 9 (SEMPER +LLM) of the essential set, with a union of 26. These genes are the partitions shown in the Venn diagram of Fig. 5e.
